# Short-term test-retest reliability of the human intrinsic functional connectome

**DOI:** 10.1101/755900

**Authors:** Leonardo Tozzi, Scott L. Fleming, Zachary D. Taylor, Cooper D. Raterink, Leanne M. Williams

## Abstract

Functional connectivity is frequently used to quantify the complex synchronous distributed fluctuations in neuronal activity derived from functional Magnetic Resonance Imaging and to generate network representations of human brain function. Such “functional connectomes” have great promise for mechanistic studies and for clinical translation. However, we do not know to what extent a functional connectome is stable over time for an individual. In the present work, we evaluate the short-term test-retest reliability of functional connectomes in a large publicly available sample of healthy participants (N=833) scanned on two consecutive days. We also assess the consequences on reliability of three methodological procedures for which a clear guideline in the community is lacking: atlas choice, global signal regression and thresholding. By adopting the intraclass correlation coefficient as a reliability metric, we demonstrate that a relatively small portion of the intrinsic functional connectome is characterized by good (4-6%) to excellent (0.08-1%) stability over time. In particular, connectivity between prefrontal, parietal and temporal areas appears to be especially stable over short timescales. Also, while unreliable edges of the functional connectome are generally weak in terms of average functional connectivity, reliable edges are not necessarily strong. Methodologically, we demonstrate that multimodal parcellation and averaging of connections within known networks are practices that improve reliability. Harnessing this knowledge, for example by honing in on the reliable portion of the connectome, offers one way forward for studies of trait-like features within the normative connectome and for discovery of biomarkers in clinical cohorts.

## 1 Introduction

The human brain is an extraordinarily complex network comprising one hundred billion neurons, each connected to an average of 7,000 other neurons. This yields between 100 trillion and 1 quadrillion synapses, depending on a person’s age [1]. Current research in neuroscience suggests that it is the architecture and dynamic interactions of neurons that give rise to complex phenomena, such as cognition and emotion [2, 3, 4]. This has been called the “functional connectome” and over the past three decades, several studies have characterized it in vivo (see for example [5]). However, we do not yet know to what extent the functional connectome is stable over time for an individual. In the present work, our aim is to explore the short-term reliability (in the order of days) of functional connectomes. We believe this is a fundamental step towards the identification of a persistent representation of brain function, which will facilitate the mapping of cognitive processes and will be critical for linking connectivity disruptions to brain disorders.

Over the past three decades, functional connectivity (FC) has become a well-established approach to measuring the functional connectome by using functional Magnetic Resonance Imaging (fMRI). The term FC refers to synchronous distributed fluctuations in neuronal activity and is thought to represent a correlate of the dynamic interaction between neurons located in different brain areas [6]. In most studies, FC is measured by computing a Pearson correlation of the fMRI derived Blood-Oxygen-Level-Dependent (BOLD) timeseries of a set of regions while the participant is awake and not performing any task [7]. This is what we will intend as FC in the present work, but it is important to note that FC can also be computed while participants are not completely idle and using a wide array of methods besides correlations, such as independent component analysis, analyses in the frequency domain, Bayesian models and dynamic approaches (for review, [7, 8, 9, 10]).

In recent years, FC has been used to answer important questions about both healthy and disordered brain function. For example, Psychiatry and Neurology have turned to FC to develop new diagnostic tests, predict treatment response and relate brain function to symptoms [11, 12]. This is in answer to an urgent need for quantitative correlates of brain illnesses and network-based approaches show great promise to this end [13]. However, when testing candidate measures for clinical applications, it is important to consider that measures of robust group-level effects, which make up a large portion of the existent literature, might not necessarily be suited to make inferences about individuals [14]. For example, a network might show consistent FC values because of its low between-subjects variability, but this same characteristic would make it unsuitable to investigate its correlations with measures that might be highly variable between subjects (symptoms, personality, etc.). Also, one reasonable characteristic of a measure considered for clinical applications is repeatability: the same test on the same individual after a short period of time should return very similar results [15]. Therefore, the combination of high between-subjects variability, coupled with low within-subject variability, indicates reliability of a measure and is advantageous when trying to relate biomarkers to individual traits.

To date, only a relatively small number of studies has examined in detail the reliability of the functional connectome. There is consensus that connectomes tend to become more reliable the longer the duration of the scan [16, 17, 18]. However, which and how many functional connections can be measured reliably is unclear. Some studies, for example, report that functional connections have only “fair” reliability on average (as defined by [19]) but others report “good” or “excellent” reliability for large (>25%) portions of the functional connectome, in particular of well-characterized functional networks such as the default mode, frontoparietal and dorsal attention networks [16, 17, 20]. Reliability of global and local graph metrics derived from functional connectomes has also been shown to only be in the “fair” range, but can still be considered statistically significant [18]. One reason for these discrepancies might be that most studies only used relatively small samples (N ≤ 50) and had long intervals between scanning sessions (days or months). Sample size has been found to affect reliability estimates [18] and shorter time intervals are better suited for measuring test-retest reliability, since they minimize the variability introduced by ancillary factors [15]. Also, there are several choices that are routinely made when computing functional connectomes and it is unclear how these might affect their reliability. Examining all such possible choices is beyond the possibilities of the current work, but we focus here on three procedures for which a clear guideline in the community is lacking. The first is the set of regions (atlas) used to compute FC. Choosing atlases with a higher number of regions has been shown to increase reliability [18], but it is unclear if a consensual pattern of reliable connections exists across atlases. Second, global signal regression (GSR) is often used as a denoising procedure but potentially affects FC and has been suggested to decrease reliability [17, 21, 22]. Third, to compute graph metrics based on the functional connectome that require sparsity, usually weaker connections are removed, either based on an absolute or relative threshold, i.e. edges below a certain value or the bottom nth percentile. Different graph metrics appear to have highest reliability at different thresholds [18]. It is, however, unclear if edge strength and reliability are related, how thresholding affects the reliability of edges, and even if the same edges are retained when functional connectomes from the same individual are thresholded independently.

In the present work, we explore short-term test-retest reliability of functional connectomes in a very large sample of healthy individuals scanned on two consecutive days. We leverage the entire Human Connectome Project (HCP) Healthy Young Adult data release, so that, to the best of our knowledge, our reliability analysis of FC is the largest ever conducted. This dataset makes use of cutting-edge acquisition and preprocessing techniques, has very long resting state sessions (30 min), is publicly available and is a widely used “gold standard” for transparent and reproducible methods testing. In these ideal conditions, first, we assess reliability of the edges and known networks that make up the functional connectome. Then, we examine the impact on reliability of atlas choice, global signal regression and thresholding.

## 2 Methods

All scripts to reproduce our analyses and plots are available on GitHub at: https://github.com/leotozzi88/repeatability_study. The data processing flow is shown in Figure 1, along with the name of the scripts in this repository corresponding to each analysis step.

**Figure 1:**
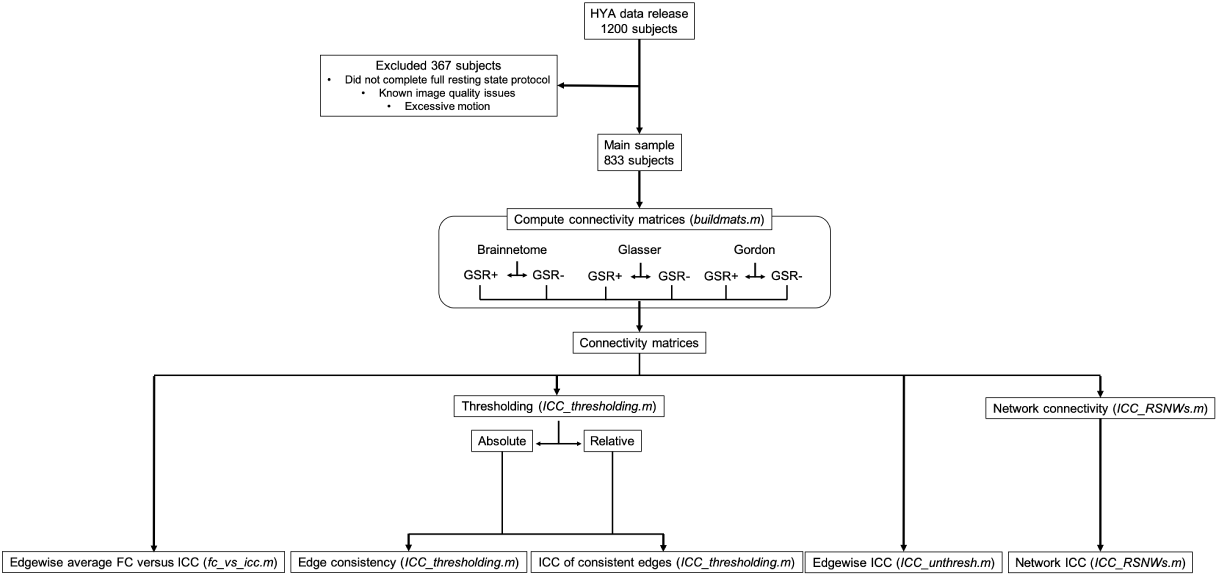
Study flow. In this diagram, the selection of the final sample and analysis steps are summarized. For each analysis step, we report the name of the github script available at https://github.com/leotozzi88/repeatability_study to reproduce the results. Abbreviations: HYA=healthy young adult, GSR-=no global signal regression, GSR+=global signal regression, ICC=intraclass correlation coefficient, FC=functional connectivity.

### 2.1 Dataset

Our sample is derived from the HCP Healthy Young Adult release, a large public dataset of 1200 subjects aged between 22 and 35 years without any psychiatric or neurological disorder [23]. The acquisition parameters and minimal preprocessing of these data are described in [24]. Briefly, participants underwent a large number of MRI scans, that included T1 and T2 weighted structural imaging, diffusion tensor imaging, and nearly 2 hours of resting-state and task multiband fMRI. For the present study, we used one hour of resting-state fMRI collected on each participant during four 15-min scans (1200 time-points each, 2 runs acquired with RL phase encoding and 2 with LR) split in two scanning sessions over two days.

To select our sample, we accessed the data at https://db.humanconnectome.org. Using the online filtering options, we selected only participants who had completed the full resting state scanning protocol and had no known quality issues. This returned a total of 860 subjects, each with four resting state fMRI runs. For these, we downloaded the framewise relative root mean square realignment estimates (RMS) and fMRI data denoised using ICA-FIX [25]. All analyses were conducted in greyordinate space, i.e. they were constrained to the grey matter by using files in the CIFTI format, thus taking full advantage of HCP preprocessing and minimizing non-neuronal signal [24].

### 2.2 Connectivity matrix construction

Each dense denoised timeseries resting state file was parcellated using connectome workbench (wb_command -cifti-parcellate) to obtain the mean timeseries in each atlas region (see below). All further analyses were conducted in Matlab R2018a (9.4.0.949201) for Mac (The MathWorks, Inc.). For each subject, parcellated timeseries as well as framewise RMS were loaded. Then, GSR was performed (see below). A high-pass filter (0.008 Hz cut-off) was applied to the timeseries. High-frequencies were retained to avoid excessive loss of degrees of freedom due to the very low TR [26]. Volumes with RMS>0.30 were flagged as containing motion and were removed from the timeseries [27]. We also excluded any subject for which volumes flagged for motion exceeded 15% in any of the 4 resting state runs. This step resulted in a final sample size of 833. The two runs within each one of the two session were then concatenated and Pearson correlation between all the timeseries was used to obtain a connectivity matrix for each session. At the end of this procedure, each subject had 6 matrices (3 atlases, with and without GSR) for each of 2 sessions, for a total of 9,996 connectivity matrices.

### 2.3 Connectivity of established resting state networks

From each connectivity matrix, the average FC within each of 12 established resting state networks was computed using the labels of the Gordon atlas [28]. These resting state networks were: default mode, parieto-occipital, fronto-parietal, salience, cingulo-opercular, medial-parietal, dorsal attention, ventral attention, visual, supplementary motor (hand), supplementary motor (mouth) and auditory. The reliability of these aggregate network statistics was then assessed.

### 2.4 Test-retest reliability

Intraclass correlation coefficient (ICC) as implemented in Matlab (https://www.mathworks.com/matlabcentral/fileexchange/22099-intraclass-correlation-coefficient-icc) was used to quantify the test-retest reliability of FC. In particular, we assessed the consistency among measurements under the fixed levels of the session factor, in line with previous work [17, 29]. This measure has been named ICC ‘C-1’ in [30] or ICC(3,1) in [31]. To calculate ICC for all our FC values, first, the FC values in the upper triangle of each subject’s connectivity matrix were entered as rows in two large matrices (one matrix for each session, one row per subject in each matrix). Then, the corresponding columns of these matrices were compared to obtain an ICC value. Since the number of features (and thus ICCs) was very large (from 21,945 to 61,776 depending on atlas), we report the median, minimum and maximum ICC as well as the portion of functional connections having poor (<0.40), fair (0.40-0.60), good (0.60-0.75) or excellent (>0.75) ICC as defined by [19]. This procedure was conducted for connectivity matrices obtained using all atlases, with and without GSR as well as for the resting state networks average FC.

### 2.5 Effects of atlas on reliability

For each subject and each session, we computed functional connectomes using three atlases that are widely used within the neuroimaging community and available in CIFTI format. The first is the Brainnettome atlas, derived by structural and functional connectivity [32]. The second is the Glasser atlas, based on the multimodal cortical parcellation of HCP participants [33]. The third is the Gordon atlas, which is based on resting state functional connectivity and provides labels identifying well-established resting state networks [28]. Since none of three atlases includes subcortical structures, these were derived from the Freesurfer segmentation [34] and added to each CIFTI dense label file using connectome workbench (wb_command -cifti-create-dense-from-template).

### 2.6 Effect of global signal regression on reliability

Immediately after loading the timeseries data in Matlab, the mean of the grayordinate timeseries from all regions was regressed from each timeseries to produce a set of GSR corrected timeseries [35]. Analyses then proceeded in the same way separately for GSR corrected (GSR+) and non-corrected (GSR-) timeseries.

### 2.7 Effect of edge strength on reliability

To get a measure of edge strength, we averaged the FC of all edges across the 2 sessions and across all subjects. Then, for each atlas, we plotted edge strength versus the ICC of the edge calculated as outlined above.

### 2.8 Effects of thresholding on reliability

To test the effects of thresholding the functional connectome on reliability estimates and if the same edges would be consistently retained across sessions, we proceeded as follows. First, we defined 20 evenly spaced threshold values from 0.05 to 1. For each value and each connectivity matrix, 2 new matrices were created using functions from the brain connectivity toolbox [36]. In the first matrix, all FC values below the threshold were set to 0 (absolute threshold). In the second, the proportion of strongest FC values corresponding to the threshold was retained (relative threshold). Then, for each absolute and relative threshold, we examined all edges that were retained at least in one session. We computed the ratio between the number of participants in which each edge was retained at both timepoints versus the ones in which it was retained at least once. This quantity, which we name “ratio of consistent edges”, or “consistency ratio” for short, goes from 0 (each time the edge is retained, it is only retained in one session) to 1 (each time the edge is retained, it is retained in both sessions). We report the median, minimum and maximum ratio of consistent edges for each threshold and atlas as well as the proportion of poor (<0.40), fair (0.40-0.60), good (0.60-0.75) or excellent (>0.75) edges using the same cut-offs as for ICC for convenience [19]. Finally, for each absolute and relative threshold, we computed the ICC as described above only in subjects for which the edge was retained in both sessions, so as not to bias our calculation by the edge potentially being set to 0 because of thresholding in one session.

### 2.9 Confirmation of results in non-related sub-sample

Since several subjects in the Healthy Young Adult dataset share family membership and a significant portion of variance in FC is explained by genetics [17, 37, 38], we reran our analyses on the unrelated participants in our dataset to confirm our results (N=83).

## 3 Results

### 3.1 Test-retest reliability of the functional connectome

Regardless of atlas and without performing GSR (see below), the majority of edges of the functional connectome were in the “fair” reliability range (Figure 2). Median ICC ranged from 0.41 to 0.47 depending on atlas (Table 1). When examining the average ICC of all edges touching each node, the subgenual ACC and inferior temporal lobe had the lowest average reliability. Average ICC was also low in areas immediately adjacent to the corpus callosum (cingulate cortex). The most reliable nodes on average were located in the superior parietal and middle temporal lobes (Figure 3). “Good” edges always represented a relatively small portion compared to the total (6-8%) but still numbered in the thousands. These connections mostly involved the inferior parietal and middle temporal lobes, but were also present in the prefrontal, superior parietal and occipital lobes (Figure 4). “Excellent” edges were relatively rare and consistently less than a hundred (0.08-0.10% of total connections). These were mostly intra-hemispherical and predominantly connected the prefrontal, parietal and temporal lobes (Figure 5).

**Figure 2:**
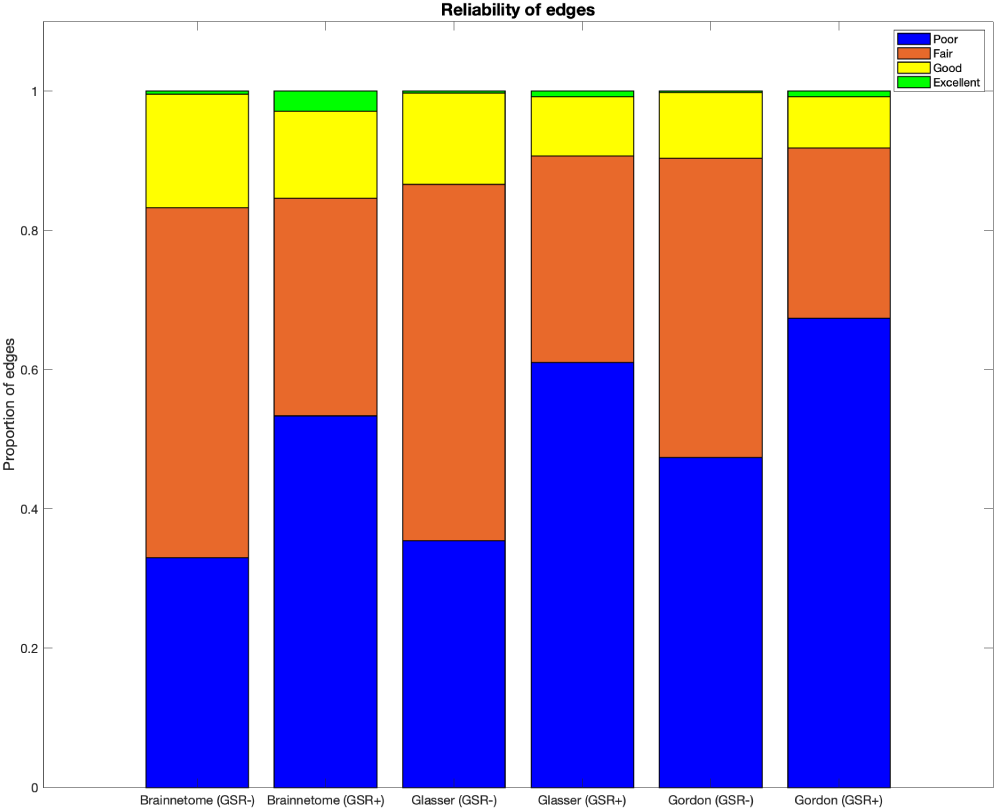
Reliability of functional connectome edges. For the Brainnetome, Glasser and Gordon atlases with and without performing GSR we show the proportion of edges having poor (ICC<0.40), fair (ICC=0.40-0.60), good (ICC=0.60-0.75) or excellent (ICC>0.75) reliability, defined in accordance to [19]. Abbreviations: ICC=intraclass correlation coefficient, GSR-=no global signal regression, GSR+=global signal regression.

**Table 1:**
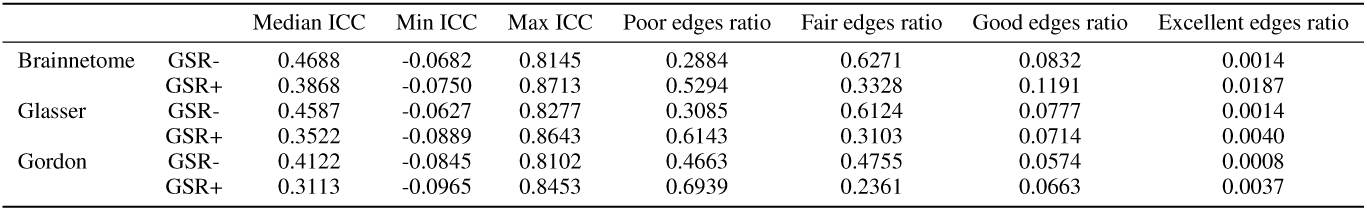
Reliability of edges in the functional connectome. We show the median, minimum and maximum ICC of functional connectomes computed using three different atlases, with or without global signal regression. We also show the proportion of edges having poor (ICC<0.40), fair (ICC=0.40-0.60), good (ICC=0.60-0.75) or excellent (ICC>0.75) reliability, defined in accordance to 20. Abbreviations: ICC=intraclass correlation coefficient, GSR-=no global signal regression, GSR+=global signal regression.

|  |  | Median ICC | Min ICC | Max ICC | Poor edges ratio | Fair edges ratio | Good edges ratio | Excellent edges ratio |
| --- | --- | --- | --- | --- | --- | --- | --- | --- |
| Brainnetome | GSR- | 0.4688 | -0.0682 | 0.8145 | 0.2884 | 0.6271 | 0.0832 | 0.0014 |
|  | GSR+ | 0.3868 | -0.0750 | 0.8713 | 0.5294 | 0.3328 | 0.1191 | 0.0187 |
| Glasser | GSR- | 0.4587 | -0.0627 | 0.8277 | 0.3085 | 0.6124 | 0.0777 | 0.0014 |
|  | GSR+ | 0.3522 | -0.0889 | 0.8643 | 0.6143 | 0.3103 | 0.0714 | 0.0040 |
| Gordon | GSR- | 0.4122 | -0.0845 | 0.8102 | 0.4663 | 0.4755 | 0.0574 | 0.0008 |
|  | GSR+ | 0.3113 | -0.0965 | 0.8453 | 0.6939 | 0.2361 | 0.0663 | 0.0037 |

**Figure 3:**
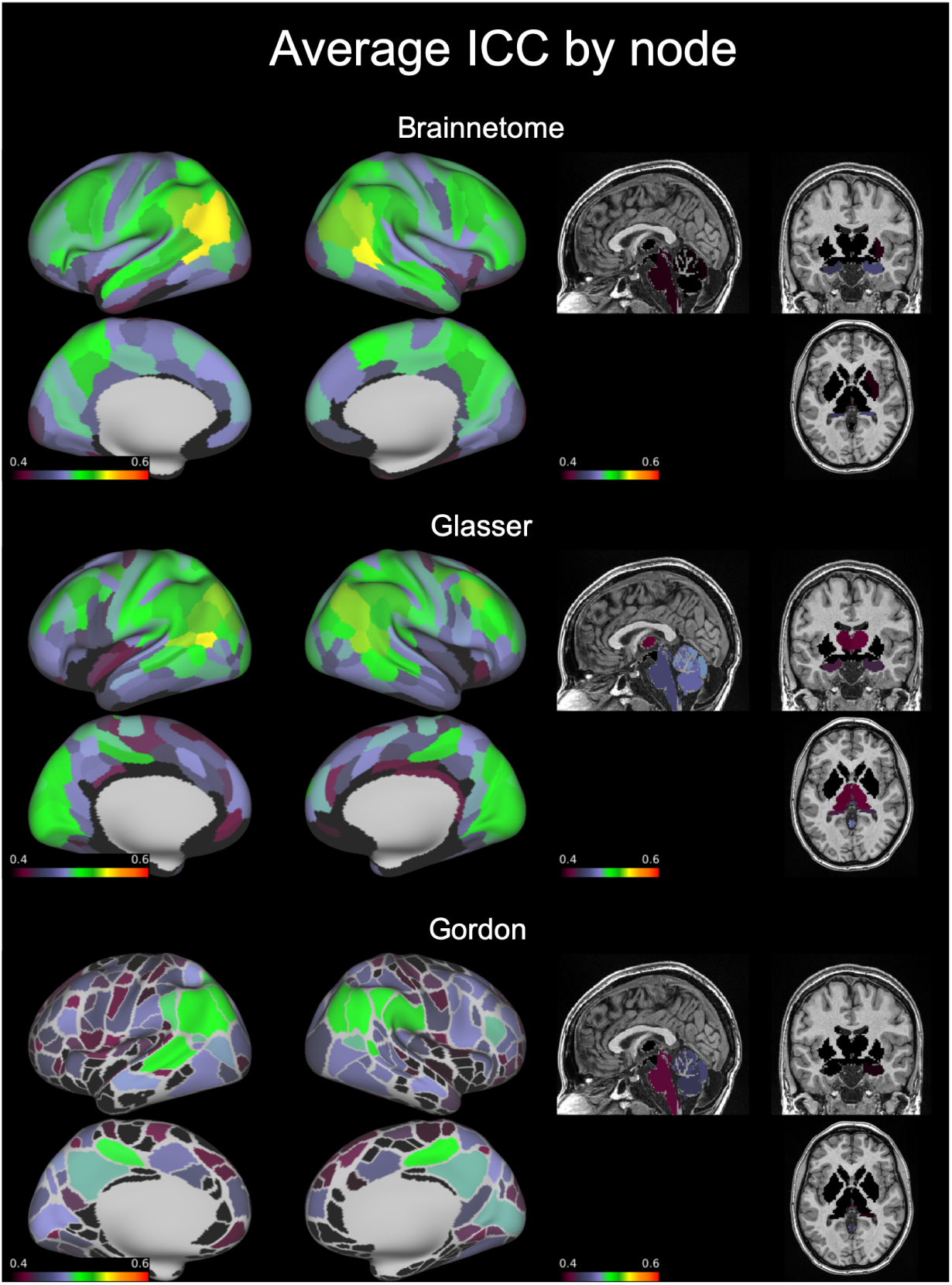
Average ICC of functional connections by node. On an inflated brain we show, for each node of the Brainnetome, Glasser and Gordon atlases, the average ICC of all functional connections involving the node. Abbreviations: ICC=intraclass correlation coefficient.

**Figure 4:**
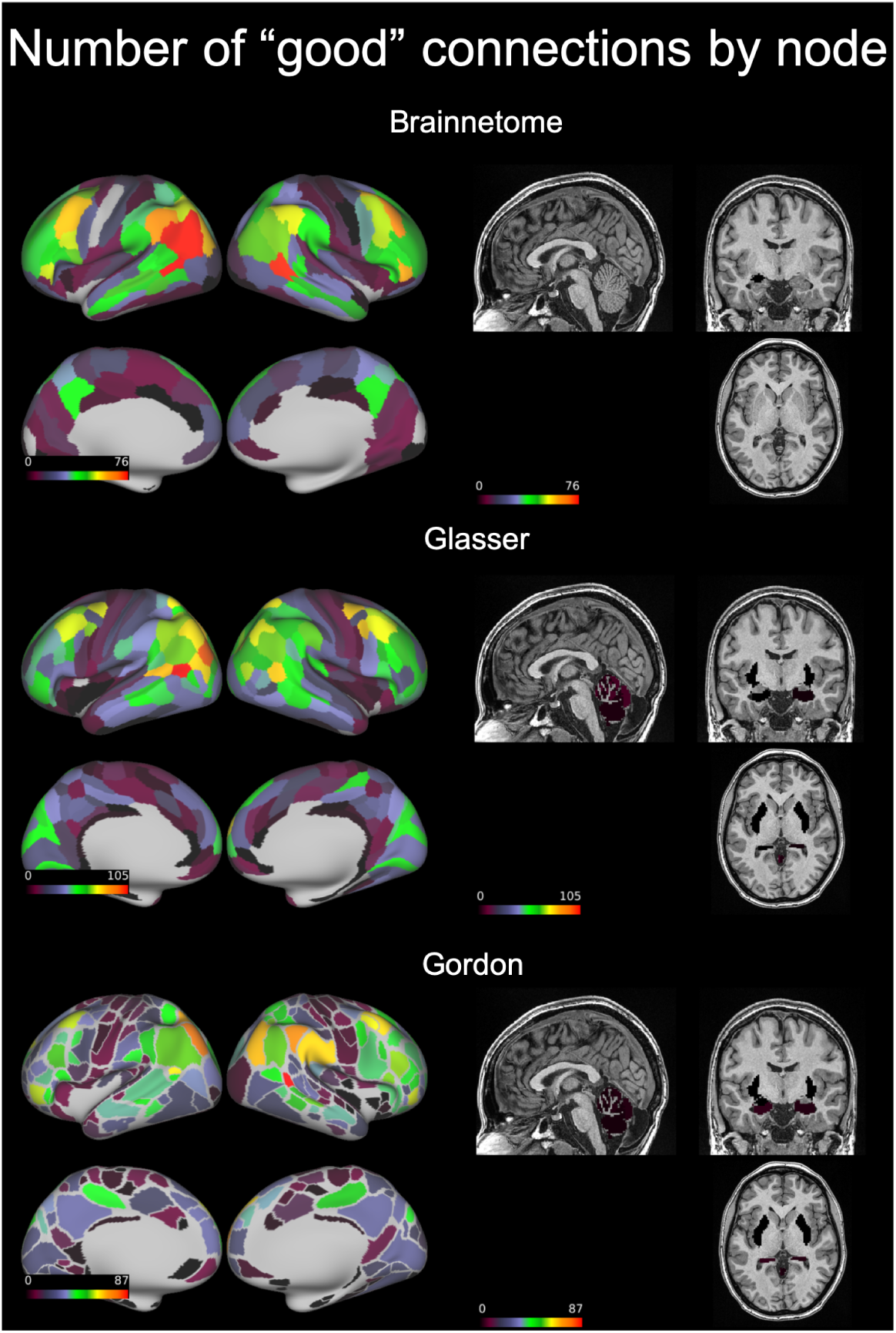
Number of functional connections with “good” reliability by node. On an inflated brain we show, for each node of the Brainnetome, Glasser and Gordon atlases, the number of functional connections involving the node with intraclass correlation coefficient=0.60-0.75.

**Figure 5:**
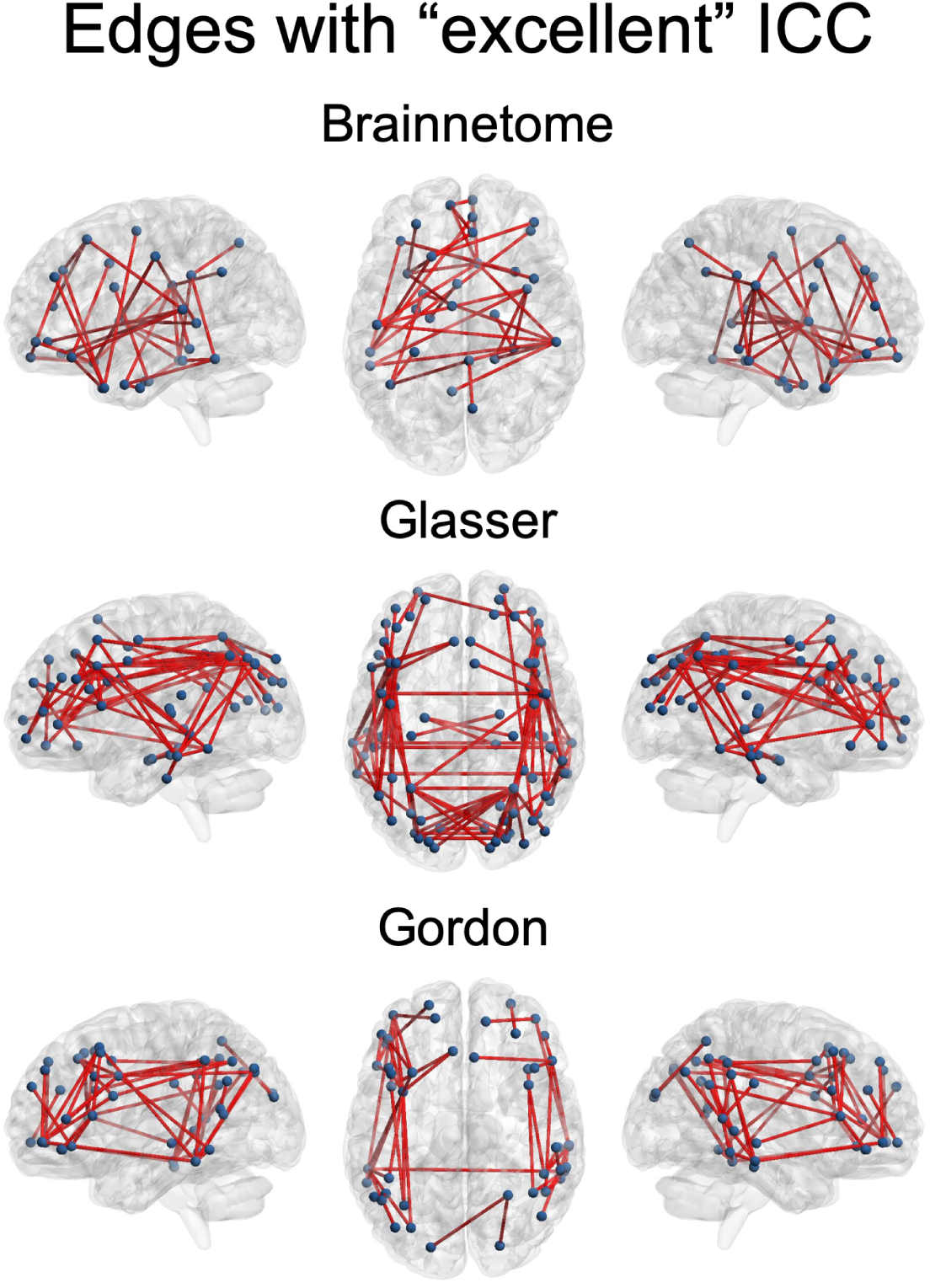
Connections with “excellent” reliability. On a transparent brain we show the connections having “excellent” reliability (intraclass correlation coefficient>0.75) in the Brainnetome, Glasser and Gordon atlases. These consistently involved the superior parietal and middle temporal lobes as well as the dorsolateral prefrontal cortex. Abbreviations: ICC=intraclass correlation coefficient.

### 3.2 Test-retest reliability of resting state networks

Average FC within established resting state networks always had “good” reliability. The most reliable network was the parieto-occipital (ICC=0.73), followed by the medial-parietal (ICC=0.71) and auditory (ICC=0.71). The least reliable were the salience (ICC=0.63), dorsal attention (ICC=0.64) and supplementary motor (mouth) (ICC=0.64) (Figure 6, Table 2).

**Figure 6:**
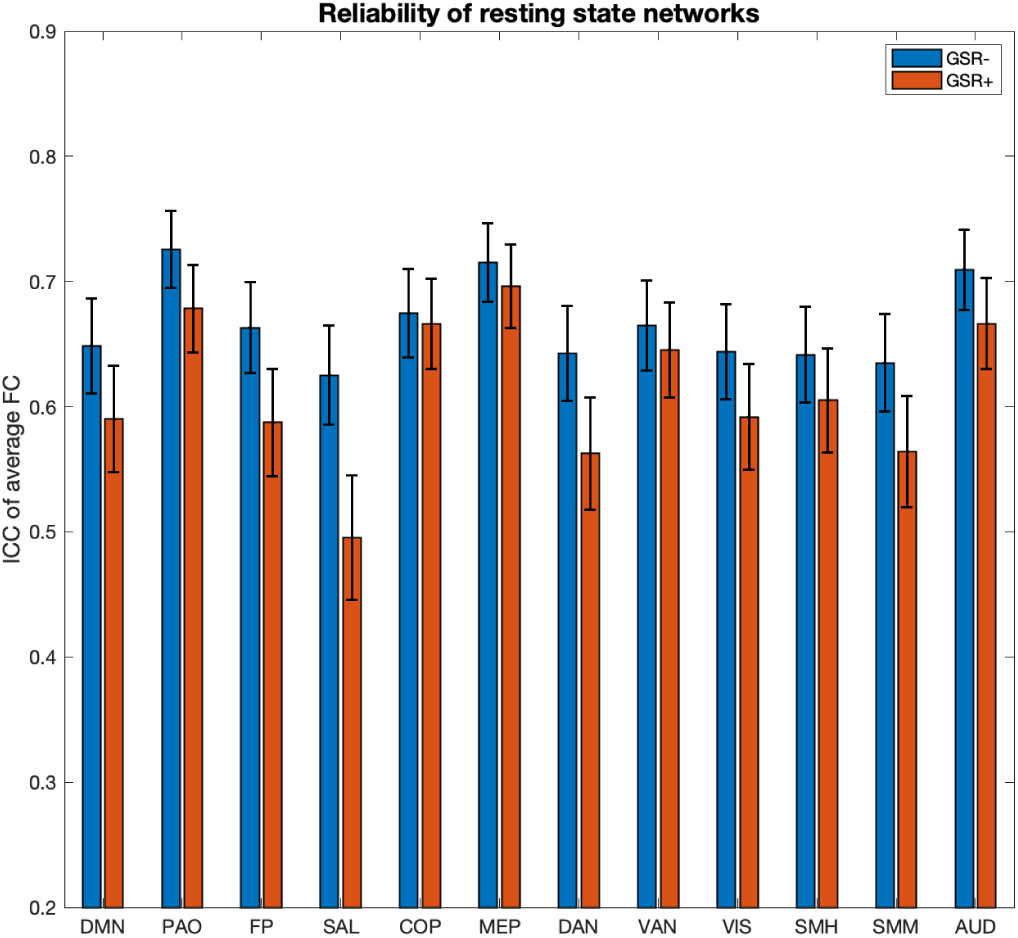
Reliability of known resting state networks. We show the ICC and confidence intervals for the average connectivity within known resting state networks defined in accordance to [19]. Abbreviations: ICC=intraclass correlation coefficient, GSR-=no global signal regression, GSR+=global signal regression, DMN=default mode network, PAO=parieto-occipital, FP=fronto-parietal, SAL=salience, COP=cingulo-opercular, MEP=medial parietal, DAN=dorsal attention network, VAN=ventral attention network, VIS=visual, SMH=supplementary motor (hand), SMM=supplementary motor (mouth), AUD=auditory.

**Table 2:** Reliability of known resting state networks. We show the ICC and confidence intervals for the average connectivity within known resting state networks defined in accordance to [28]. Abbreviations: ICC=intraclass correlation coefficient, GSR-=no global signal regression, GSR+=global signal regression, CI=confidence interval, DMN=default mode network, PAO=parieto-occipital, FP=fronto-parietal, SAL=salience, COP=cingulo-opercular, MEP=medial parietal, DAN=dorsal attention network, VAN=ventral attention network, VIS=visual, SMH=supplementary motor (hand), SMM=supplementary motor (mouth), AUD=auditory.

|  | ICC |  | Lower CI |  | Upper CI |  |
| --- | --- | --- | --- | --- | --- | --- |
|  | GSR- | GSR+ | GSR- | GSR+ | GSR- | GSR+ |
| DMN | 0.6485 | 0.5900 | 0.6862 | 0.6326 | 0.6074 | 0.5439 |
| PAO | 0.7256 | 0.6783 | 0.7562 | 0.7133 | 0.6918 | 0.6398 |
| FP | 0.6631 | 0.5873 | 0.6995 | 0.6301 | 0.6233 | 0.5410 |
| SAL | 0.6252 | 0.4953 | 0.6648 | 0.5449 | 0.5820 | 0.4423 |
| COP | 0.6744 | 0.6661 | 0.7098 | 0.7022 | 0.6356 | 0.6265 |
| MEP | 0.7153 | 0.6960 | 0.7469 | 0.7294 | 0.6804 | 0.6593 |
| DAN | 0.6426 | 0.5625 | 0.6808 | 0.6072 | 0.6010 | 0.5142 |
| VAN | 0.6648 | 0.6454 | 0.7011 | 0.6834 | 0.6251 | 0.6040 |
| VIS | 0.6437 | 0.5918 | 0.6818 | 0.6342 | 0.6022 | 0.5458 |
| SMH | 0.6413 | 0.6050 | 0.6796 | 0.6464 | 0.5996 | 0.5601 |
| SMM | 0.6350 | 0.5640 | 0.6739 | 0.6086 | 0.5927 | 0.5159 |
| AUD | 0.7095 | 0.6664 | 0.7417 | 0.7025 | 0.6741 | 0.6269 |

### 3.3 Effects of atlas on reliability

The Brainnetome and Glasser atlases had comparable median ICCs (0.47 and 0.46 respectively, Table 1). The proportion of “good” and “excellent” edges was also comparable (8% and 0.14% for both atlases, respectively). However, since the Glasser atlas has more nodes than the Brainnetome (229 versus 379), it generated a much larger number of edges (71,631 versus 26,106). Therefore, using the Glasser atlas returned a higher absolute number of “good” and “excellent” edges (Figure 5). Median ICC for the Gordon atlas (352 nodes) was lower (0.41), as well as the proportion of “good” (5%) and “excellent” (0.08%) edges.

### 3.4 Effect of global signal regression on reliability

Regardless of atlas, performing GSR led the reliability of most edges to go from “fair” to “poor”. In particular, the ratio of “poor” edges increased from 29% to 53% in the Brainnetome atlas, 31% to 61% in the Glasser and from 47% to 69% in the Gordon. Conversely, the ratio of good edges decreased from 63% to 33%, from 61% to 31% and from 47% to 24% respectively. The ratio of “good” edges, however, increased for the Brainnetome and Gordon atlases (from 8% to 12% and from 6% to 7% respectively). Also, GSR increased the ratio of “excellent” edges in all atlases: from 0.1% to 2% in the Brainnetome, from 0.1% to 0.4% in the Glasser, from 0.08% to 0.4% in the Gordon (Table 1 Figure 2). GSR also reduced the reliability of average FC of resting state networks (Table 2, Figure 6).

### 3.5 Effect of edge strength on reliability

Overall, the distribution of edges in the functional connectome had its highest density in an interval of r=0.10-0.30 and a corresponding reliability ranging from “poor” to “fair” (ICC=0.30-0.50) (Figure 7). The FC of edges with “excellent” reliability varied across a wide range for each of the atlases: Brainnetome, r=0.25-0.61, median=0.40; Glasser r=0.12-0.61, median=0.34; Gordon r=0.13-0.44, median=0.22. The range was even broader for edges with “good” reliability (respectively r=-0.06-0.85, median=0.34; r=-0.04-0.80, median=0.25; r=-0.01-0.80 median=0.21) as well as “fair” reliability (respectively r=-0.13-0.83, median=0.20; r=-0.08-0.72, median=0.10; r=-0.05-0.60 median=0.10). Edges with poor reliability (ICC<0.40) tended to have lower FC in all three atlases: respectively r=-0.04-0.45, median=0.08; r=-0.04-0.23, median=0.04; r=-0.03-0.22, median=0.03.

**Figure 7:**
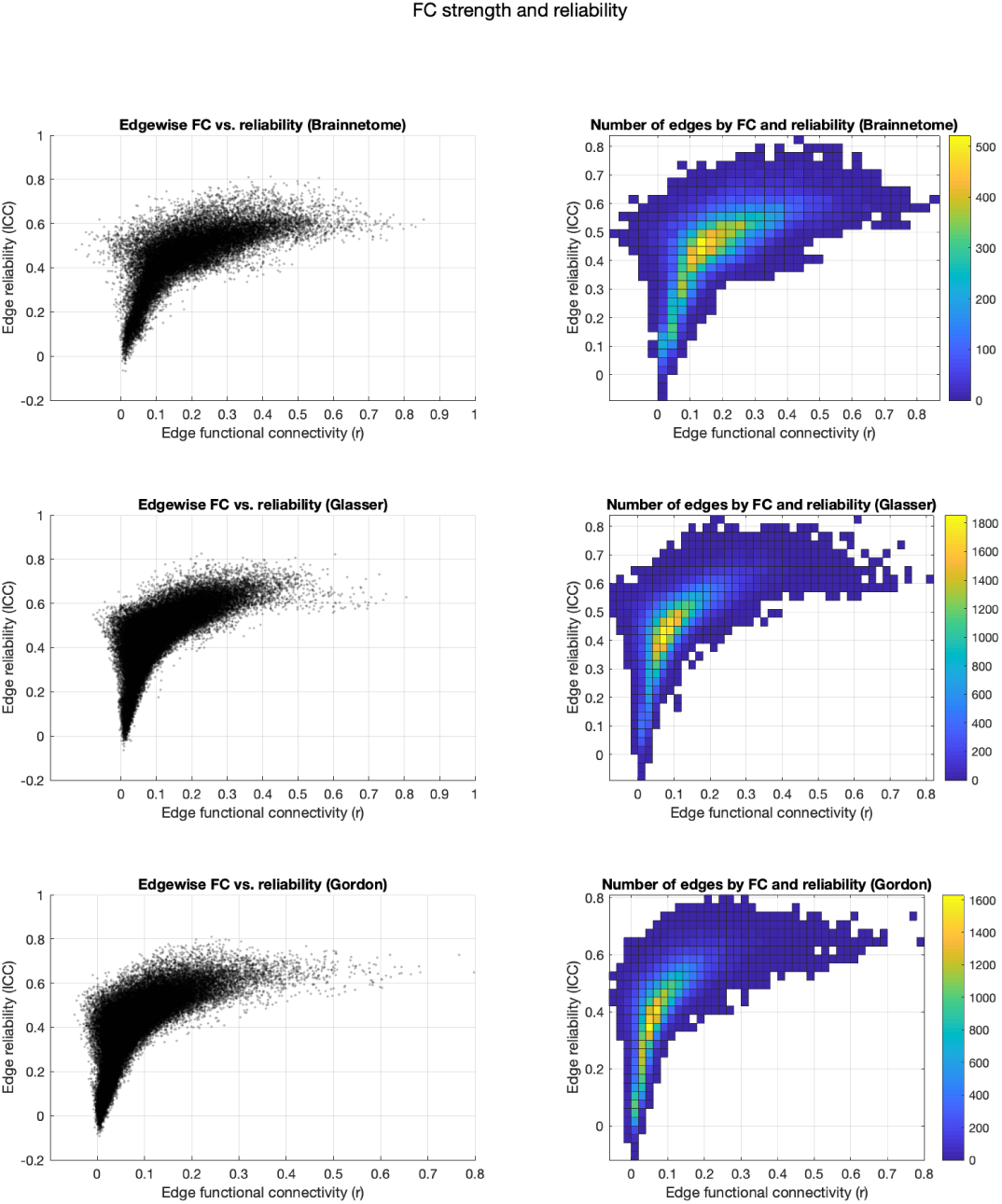
Effects of edge strength on reliability. Plotting the strength of each edge of the Brainnetome, Glasser and Gordon atlases versus their ICC (left) showed that edges with higher FC were mostly in the “fair” reliability range (ICC=0.40-0.60). FC of edges with higher reliability (ICC>0.70) was mostly between r=0.10 and r=0.70 and edges with very low reliability (ICC<0.30) also had low FC (r<0.20). The distribution of edges (right) had its highest density in an interval of r=0.10-0.30 and a corresponding reliability ranging from “poor” to “fair” (ICC=0.30-0.50). Abbreviations: ICC=intraclass correlation coefficient, FC=functional connectivity, r=Pearson correlation coefficient.

### 3.6 Effects of thresholding on reliability

The median rate of consistent edges decreased as thresholds (both absolute and relative) became more stringent. The effect of absolute thresholding was a sharp decline in the median ratio of consistent edges, such that from r>0.45 upwards, edges with “excellent” consistency ratio were only around 1% of those retained at least once. Using a relative threshold, the median ratio of consistent edges showed a linear decline as the thresholding became more stringent, mirrored by a proportional increase of edges demonstrating a “poor” consistency ratio (Tables S1-S3, Figure 8). For edges retained at both timepoints, median ICC decreased as a result of increasingly stringent absolute thresholds and slightly increased (∼0.10) with increasingly stringent relative thresholds. ICC of most edges being consistently retained after absolute thresholding was “poor”. After relative thresholding, on the other hand, the largest proportion of consistently retained edges was “fairly” reliable. Strict relative thresholds saw a slight increase in consistently retained edges that had “excellent” reliability, mirrored by a decline in “fair” ones (Table S4-S6, Figure 9).

**Figure 8:**
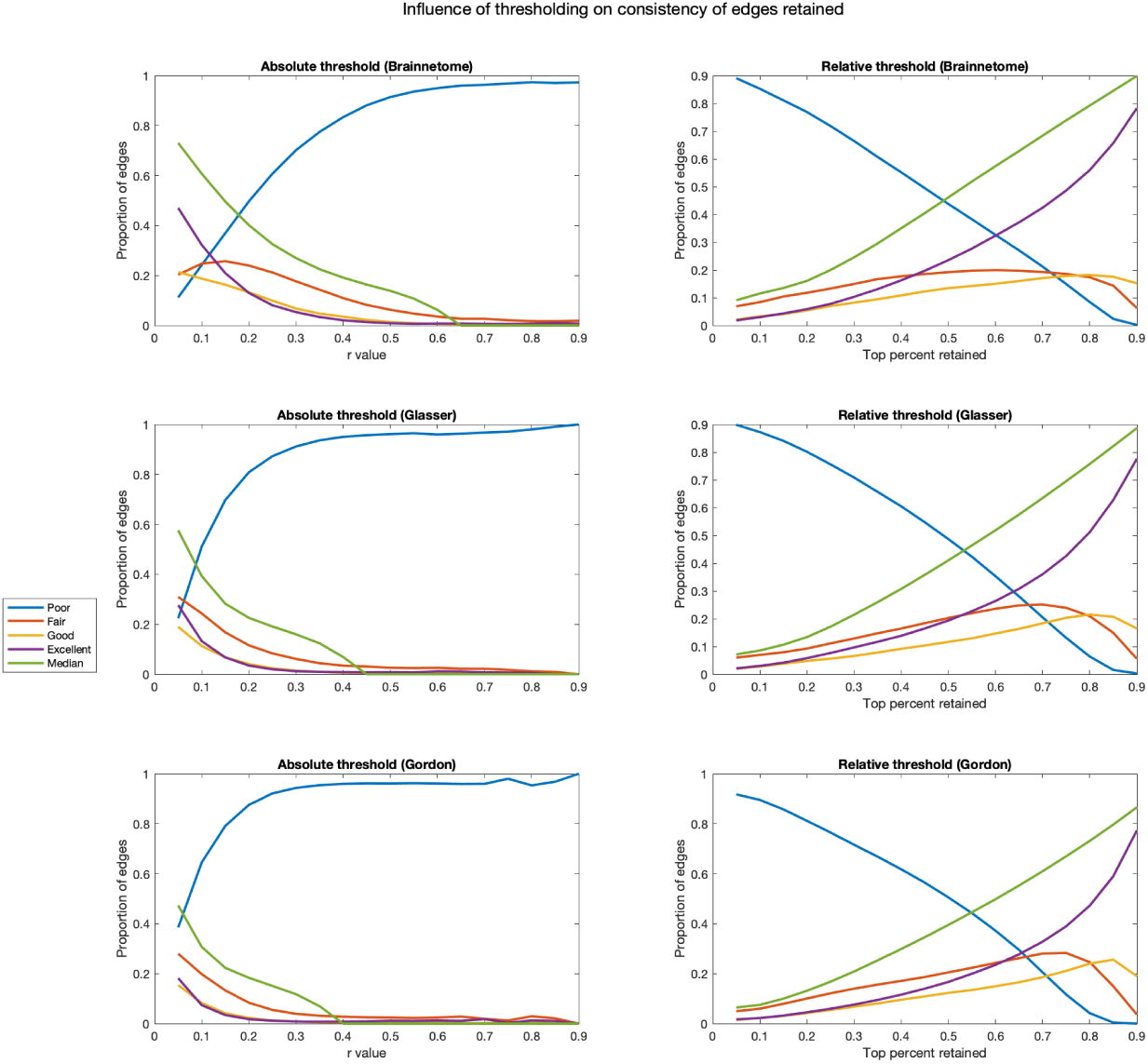
Effects of thresholding on edge retention. In the Brainnetome, Glasser and Gordon atlases, for each absolute and relative threshold we show the proportion of edges that are consistently retained. As a measure of consistency, we use the number of participants in which the edge was retained at both timepoints divided by the ones in which it was retained at least once. For convenience, we then use the values defined in [19] to plot the ratio of edges having poor (ratio<0.40), fair (ratio=0.40-0.60), good (ratio=0.60-0.75) or excellent (ratio >0.75) consistency. For absolute thresholds (left) all edges below the value are set to 0, for relative ones (right) only the top percent corresponding to the threshold is retained. Abbreviations: r=Pearson correlation coefficient.

**Figure 9:**
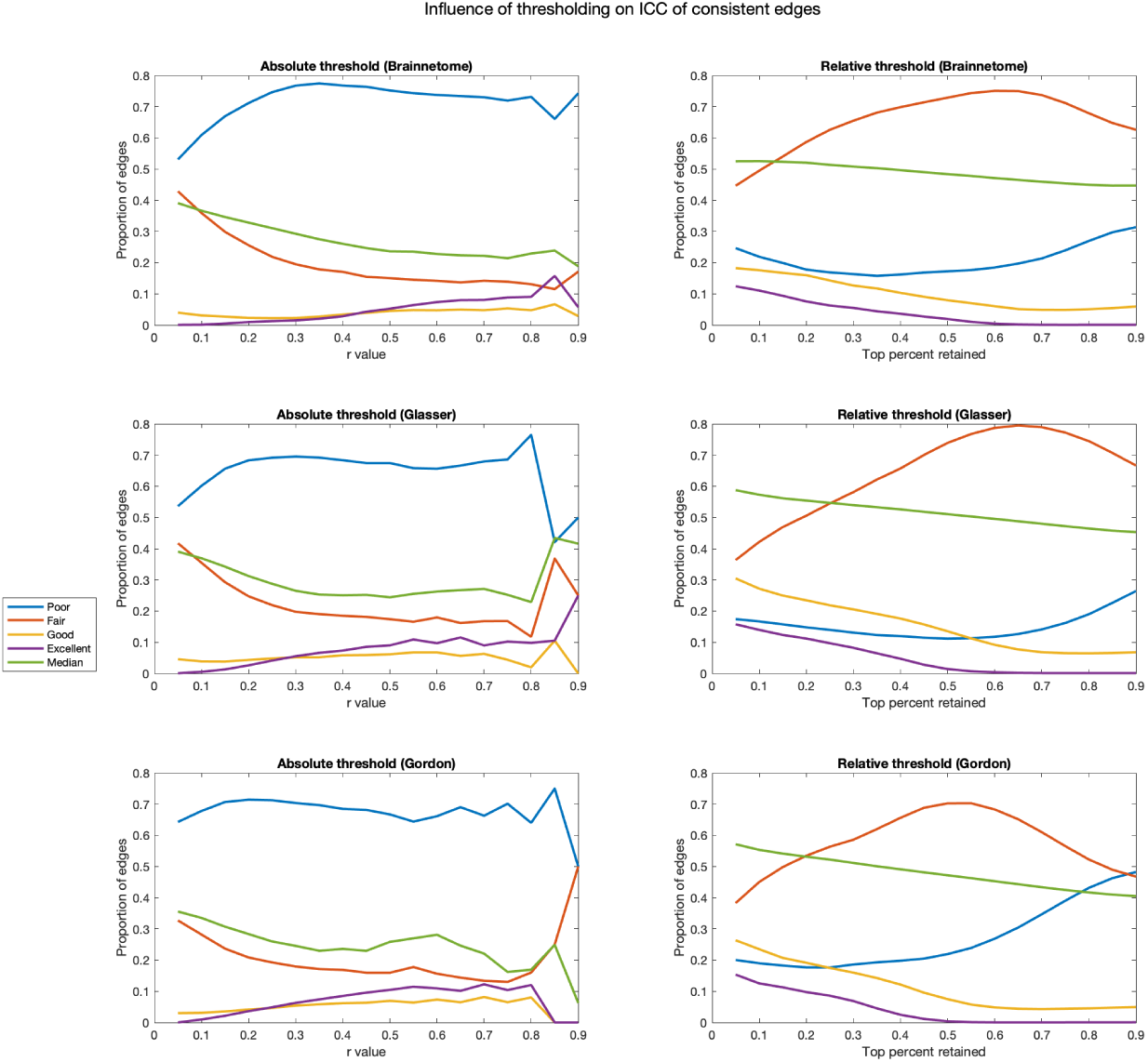
Effects of thresholding on reliability. In the Brainnetome, Glasser and Gordon atlases, for each absolute and relative threshold we show the proportion of edges having poor (ICC<0.40), fair (ICC=0.40-0.60), good (ICC=0.60-0.75) or excellent (ICC>0.75) reliability. In this calculation, only subjects for which the edge was retained in both sessions were considered. For absolute thresholds (left) all edges below the value are set to 0, for relative ones (right) only the top percent corresponding to the threshold is retained. Abbreviations: r=Pearson correlation coefficient, ICC=intraclass correlation coefficient.

### 3.7 Confirmation of results in non-related sub-sample

In a smaller sub-sample of unrelated participants, without performing GSR, the majority of edges still had “fair” reliability. The proportion of “good” and “excellent” edges was higher in all atlases (Table S7, Figure S1). When examining resting state networks, ICC was lower for the default mode, parieto-occipital, fronto-parietal, salience, cingulo-opercular, medial-parietal, dorsal attention and ventral attention networks. It was higher for the visual, supplementary motor (hand and mouth) and auditory networks. Most networks had still “good” reliability, but the default mode, fronto-parietal, salience, dorsal and ventral attention only had “fair” reliability (Table S8, Figure S2). The effects of GSR were similar to the ones observed in the full sample: it greatly increased the proportion of “poor” edges, but also caused a small increase in “good” and “excellent” ones. For resting state networks, GSR increased ICC of the default mode, parieto-occipital, fronto-parietal, medial-parietal, ventral attention networks, but decreased ICC of the others (Tables S7-S8, Figure S2). The effects of thresholding were also confirmed in the unrelated sub-sample. We observed a decrease of consistency of edges retained with increasingly strict thresholds (Table S9-S11, Figure S3). The median ICC of consistently retained edges was not strongly impacted by thresholding, but strict relative thresholds saw an increase in the proportion of “excellent” edges and a decrease of “fair” ones (Table S12-S14, Figure S4).

## 4 Discussion

In this work, we conducted the largest investigation of the short-term test-retest reliability of functional connectivity to date. We conclude that a small fraction of the intrinsic connectome has good to excellent test-retest reliability. In particular, connectivity between prefrontal, parietal and temporal areas appears to be especially stable over short timescales. Also, while unreliable edges are generally weak in terms of average functional connectivity, the most reliable edges are not necessarily strong. Methodologically, we demonstrate that multimodal parcellation and averaging of connections within known networks are practices that improve reliability.

First of all, our results provide theoretical insights about the variability in the functional architecture of the human brain. We show that, regardless of atlas, reliability of the vast majority of edges of the functional connectome is only “fair”. Still, thousands of connections had “good” reliability, and less than a hundred were “excellent”. This is in contrast with a previous study, which reported that the “good” and “excellent” edges combined were more than 25% of the functional connectome when considering 30 minutes of resting state [17]. This discrepancy might be due to the much lower sample size of the previous study (N=33), which might not have had sufficient power to detect low ICC values. Indeed, with only 2 observations per subject, for example, at least 152 subjects are needed to detect an ICC=0.20 with power = 80% and alpha = 0.05 [39]. Importantly, when considering the distribution of the most reliable edges in the brain, some consistent patterns emerged across all atlases. Not surprisingly, connections involving areas prone to susceptibility artifacts, like the subgenual anterior cingulate cortex and inferior temporal lobe had the lowest average reliability, probably because of lower signal to noise ratio. Secondly, the most reliable connections involved nodes in the superior parietal, middle temporal lobes and dorsolateral prefrontal cortex. This pattern bears a striking resemblance to what has been labeled the fronto-parietal network [40]. This network is implicated in cognitive control and interacts with other networks implicated in this function, such as the cingulo-opercular, and salience networks [41, 42]. It undergoes functional changes through development, increasing in both strength and flexibility from childhood to adulthood [43]. Crucially, topography of the fronto-parietal network is highly variable between individuals, which might explain the very high ICC we detected for these connections [44]. Interestingly, the average connectivity of the fronto-parietal network, as defined by [28], had only “good” reliability, despite presence of several “excellent” edges in this network as outlined above, suggesting that averaging connections might reduce the impact of the relatively rare “excellent” ones.

Our work informs future studies wanting to focus on portions of the functional connectome that combine high between-subjects reliability with low within-subjects reliability, such as those investigating potential clinical biomarkers or correlations with behavioral measures or self-reports. In particular, the fronto-parietal network has also been shown to be disrupted across a wide range of psychiatric disorders [45]. This finding, combined with the excellent reliability of FC we have shown for this network, highlights that the functional connections between dorsolateral prefrontal, superior parietal and middle temporal cortices are excellent potential targets for biomarker development in Psychiatry. To aid future studies wishing to focus on specific edges depending on their reliability, we provide connectivity “masks” for each atlas in our GitHub repository, for each atlas, i.e. connectivity matrices in which each edge is labelled with its reliability (poor, fair, good or high) and connectivity matrices with the ICC values for each edge (“ICC_matrices/”). Our results also indicate that another viable complimentary approach for biomarker studies is computing average connectivity of known resting state networks using an atlas, which consistently returns values that have “good” reliability. This is in line with previous studies which showed, for example, that average connectivity of the default mode network is a good candidate measure for clinical translation [46]. This also offers a very strong reduction of the number of features (from tens of thousands to a dozen), which might be desirable, depending on the study aims.

From a methodological point of view, we were able to test the consequences on reliability of atlas choice, global signal regression and thresholding. Concerning the choice of atlas, both the Brainnetome and Glasser atlases had comparable proportion of “good” and “excellent” edges, whereas the Gordon atlas showed consistently lower reliability. Given that Brainnetome and Glasser are multimodal parcellations while the Gordon is not, this finding suggests that multimodal parcellations may provide more reliable edges compared to those based on functional data alone. The Glasser atlas also had the highest absolute number of connections, which led to identification of the highest number of “excellent” edges. Researchers wishing a higher number of reliable edges could consider this atlas, remembering however that this comes at the cost of more “poor” and “fair” edges as well. Global signal regression consistently led the majority of edges from having “fair” reliability to having “poor” reliability. Also, it consistently reduced ICC of average FC within resting state networks. This is probably due to the fact that regressing the average signal from each individual subject eliminates the between-subject variability due to differences in the signal baseline, similarly to what has been shown for contrasts between conditions in task fMRI data [47]. Interestingly, the ratio of “excellent” edges consistently increased, suggesting that some edges might benefit from GSR. Therefore, as previous studies suggested, applying GSR or not can reveal complimentary insights and the choice should be carefully made depending on the study aims [22]. Examining the role between FC strength, averaged across sessions, and reliability unveiled a complex relationship. While unreliable edges are generally weak in terms of FC, the opposite is not true: the most reliable edges are not necessarily strong and lie in a broad interval of r values (r=0.12-0.61 depending on atlas). Therefore, thresholding the functional connectome based on absolute or relative edge strength had in general detrimental consequences on reliability. First of all, applying an increasingly stringent threshold for the two sessions from each subject independently led to an increasingly large amount of connections not being consistently retained. For example, when retaining only connections with r>0.20 or higher, most of the connections that survived thresholding did so in one session but not in the other. The impact of relative thresholding was overall less dramatic compared to absolute thresholding, with the median proportion of reliably kept edges crossing 0.50 only for a threshold of 0.50. Examining the impact on ICC of the edges consistently retained after thresholding showed only a slight improvement on the median ICC and ratio of “excellent” edges for increasingly stringent relative thresholds. Also, we calculated ICC only for consistently retained edges, which we showed to be relatively rare. Overall, our results suggest that the practice of thresholding functional connectomes based on edge strength should be avoided in biomarker and correlational studies, especially in the case of absolute thresholds.

Our study is not without limitations. First of all, given that a significant portion of variance in FC is explained by genetics [17, 37, 38], our analyses on the complete Healthy Young Adult dataset might have suffered from reduced between-subjects variability and our results might have overall underestimated reliability. A recent study on the same dataset [38] concluded that genetic influence on FC is only moderate. However, performing the same analyses on the subset of unrelated participants returned results that were overall comparable, but indicated slightly higher reliability. We cannot determine whether this is indeed due to the genetic influence on FC or to the smaller number of subjects reducing our power to detect edges with low ICC values as outlined above. It is also important to mention that the ICC estimates on unrelated participants had much broader confidence intervals compared to those of our main analysis because of the lower number of participants. To be certain of removing the effect of family on reliability, a target for future studies is the analysis of very large cohorts of mostly unrelated subjects.

In conclusion, our findings offer insights into addressing the “reproducibility crisis” in the field of functional neuroimaging [48]. We demonstrate that a readily identifiable portion of the intrinsic functional connectome is characterized by good to excellent stability over time. In particular, averaging of connections within a circuit without thresholding yields a robust metric for stable functional connectivity. Fronto-temporo-parietal connections appear to be especially stable over short timescales, and thus suited to delineation of trait-like markers of both normative and disrupted cognitive functions. Harnessing this knowledge, for example by honing in on this portion of the connectome, offers one way forward for studies of trait-like features within the normative connectome and for discovery of biomarkers in clinical cohorts.

