## Supplementary material for "Short-term test-retest reliability of the human intrinsic functional connectome"

**Supplementary Tables**

Table S1

|  | Absolute | | | | | | | Relative | | | | | | |
| --- | --- | --- | --- | --- | --- | --- | --- | --- | --- | --- | --- | --- | --- | --- |
| Threshold | Median ratio | Min ratio | Max ratio | Poor edges ratio | Fair edges ratio | Good edges ratio | Excellent edges ratio | Median ratio | Min ratio | Max ratio | Poor edges ratio | Fair edges ratio | Good edges ratio | Excellent edges ratio |
| 0.050 | 0.731 | 0.052 | 1.000 | 0.112 | 0.204 | 0.214 | 0.471 | 0.091 | 0.000 | 1.000 | 0.891 | 0.069 | 0.021 | 0.018 |
| 0.100 | 0.607 | 0.000 | 1.000 | 0.242 | 0.248 | 0.188 | 0.322 | 0.115 | 0.000 | 1.000 | 0.853 | 0.084 | 0.033 | 0.030 |
| 0.150 | 0.496 | 0.000 | 1.000 | 0.370 | 0.257 | 0.163 | 0.210 | 0.135 | 0.000 | 1.000 | 0.811 | 0.104 | 0.041 | 0.043 |
| 0.200 | 0.402 | 0.000 | 1.000 | 0.497 | 0.240 | 0.133 | 0.130 | 0.161 | 0.000 | 1.000 | 0.769 | 0.118 | 0.054 | 0.059 |
| 0.250 | 0.325 | 0.000 | 1.000 | 0.607 | 0.212 | 0.099 | 0.081 | 0.200 | 0.000 | 1.000 | 0.719 | 0.133 | 0.070 | 0.078 |
| 0.300 | 0.270 | 0.000 | 1.000 | 0.702 | 0.177 | 0.068 | 0.054 | 0.246 | 0.000 | 1.000 | 0.665 | 0.150 | 0.082 | 0.103 |
| 0.350 | 0.225 | 0.000 | 1.000 | 0.775 | 0.143 | 0.048 | 0.034 | 0.296 | 0.000 | 1.000 | 0.607 | 0.167 | 0.095 | 0.131 |
| 0.400 | 0.192 | 0.000 | 1.000 | 0.834 | 0.110 | 0.035 | 0.021 | 0.350 | 0.000 | 1.000 | 0.552 | 0.177 | 0.108 | 0.163 |
| 0.450 | 0.164 | 0.000 | 1.000 | 0.882 | 0.082 | 0.022 | 0.014 | 0.405 | 0.000 | 1.000 | 0.495 | 0.185 | 0.123 | 0.197 |
| 0.500 | 0.139 | 0.000 | 1.000 | 0.914 | 0.063 | 0.014 | 0.009 | 0.461 | 0.000 | 1.000 | 0.438 | 0.192 | 0.135 | 0.235 |
| 0.550 | 0.108 | 0.000 | 1.000 | 0.936 | 0.048 | 0.009 | 0.007 | 0.519 | 0.000 | 1.000 | 0.383 | 0.198 | 0.142 | 0.277 |
| 0.600 | 0.063 | 0.000 | 1.000 | 0.950 | 0.036 | 0.006 | 0.008 | 0.574 | 0.008 | 1.000 | 0.326 | 0.200 | 0.150 | 0.324 |
| 0.650 | 0.000 | 0.000 | 1.000 | 0.960 | 0.027 | 0.005 | 0.008 | 0.629 | 0.024 | 1.000 | 0.271 | 0.198 | 0.160 | 0.372 |
| 0.700 | 0.000 | 0.000 | 1.000 | 0.963 | 0.027 | 0.003 | 0.007 | 0.684 | 0.029 | 1.000 | 0.212 | 0.193 | 0.171 | 0.424 |
| 0.750 | 0.000 | 0.000 | 1.000 | 0.969 | 0.021 | 0.004 | 0.006 | 0.739 | 0.079 | 1.000 | 0.149 | 0.186 | 0.178 | 0.486 |
| 0.800 | 0.000 | 0.000 | 1.000 | 0.974 | 0.017 | 0.002 | 0.007 | 0.793 | 0.096 | 1.000 | 0.085 | 0.174 | 0.182 | 0.560 |
| 0.850 | 0.000 | 0.000 | 1.000 | 0.971 | 0.017 | 0.002 | 0.010 | 0.846 | 0.169 | 1.000 | 0.024 | 0.143 | 0.175 | 0.657 |
| 0.900 | 0.000 | 0.000 | 1.000 | 0.973 | 0.019 | 0.000 | 0.008 | 0.899 | 0.233 | 1.000 | 0.002 | 0.063 | 0.152 | 0.783 |
| 0.950 | 0.000 | 0.000 | 0.250 | 1.000 | 0.000 | 0.000 | 0.000 | 0.953 | 0.305 | 1.000 | 0.000 | 0.007 | 0.047 | 0.946 |

**Effects of thresholding on edge retention in the Brainnetome atlas.** For each absolute and relative threshold we show the proportion of edges that are consistently retained. As a measure of consistency, we use the number of participants in which the edge was retained at both timepoints divided by the ones in which it was retained at least once. For convenience, we then use the values defined in ^20^ to compute the ratio of edges having poor (ratio<0.40), fair (ratio=0.40-0.60), good (ratio=0.60-0.75) or excellent (ratio >0.75) consistency. For absolute thresholds, all edges below the value are set to 0, for relative ones (right) only the top percent corresponding to the threshold is retained.

Table S2

|  | Absolute | | | | | | | Relative | | | | | | |
| --- | --- | --- | --- | --- | --- | --- | --- | --- | --- | --- | --- | --- | --- | --- |
| Threshold | Median ratio | Min ratio | Max ratio | Poor edges ratio | Fair edges ratio | Good edges ratio | Excellent edges ratio | Median ratio | Min ratio | Max ratio | Poor edges ratio | Fair edges ratio | Good edges ratio | Excellent edges ratio |
| 0.050 | 0.576 | 0.037 | 1.000 | 0.224 | 0.310 | 0.190 | 0.277 | 0.071 | 0.000 | 1.000 | 0.899 | 0.060 | 0.020 | 0.021 |
| 0.100 | 0.393 | 0.000 | 1.000 | 0.511 | 0.243 | 0.113 | 0.132 | 0.086 | 0.000 | 1.000 | 0.873 | 0.070 | 0.028 | 0.030 |
| 0.150 | 0.282 | 0.000 | 1.000 | 0.697 | 0.168 | 0.068 | 0.067 | 0.106 | 0.000 | 1.000 | 0.841 | 0.079 | 0.038 | 0.042 |
| 0.200 | 0.226 | 0.000 | 1.000 | 0.809 | 0.116 | 0.041 | 0.035 | 0.134 | 0.000 | 1.000 | 0.801 | 0.093 | 0.048 | 0.058 |
| 0.250 | 0.191 | 0.000 | 1.000 | 0.873 | 0.083 | 0.024 | 0.020 | 0.171 | 0.000 | 1.000 | 0.756 | 0.111 | 0.056 | 0.077 |
| 0.300 | 0.160 | 0.000 | 1.000 | 0.912 | 0.061 | 0.015 | 0.012 | 0.215 | 0.000 | 1.000 | 0.709 | 0.129 | 0.066 | 0.097 |
| 0.350 | 0.124 | 0.000 | 1.000 | 0.936 | 0.044 | 0.010 | 0.010 | 0.260 | 0.000 | 1.000 | 0.657 | 0.148 | 0.078 | 0.117 |
| 0.400 | 0.069 | 0.000 | 1.000 | 0.950 | 0.035 | 0.006 | 0.009 | 0.308 | 0.000 | 1.000 | 0.605 | 0.165 | 0.092 | 0.139 |
| 0.450 | 0.000 | 0.000 | 1.000 | 0.957 | 0.031 | 0.004 | 0.008 | 0.358 | 0.000 | 1.000 | 0.548 | 0.184 | 0.104 | 0.165 |
| 0.500 | 0.000 | 0.000 | 1.000 | 0.962 | 0.026 | 0.004 | 0.008 | 0.410 | 0.000 | 1.000 | 0.487 | 0.203 | 0.117 | 0.194 |
| 0.550 | 0.000 | 0.000 | 1.000 | 0.965 | 0.025 | 0.003 | 0.007 | 0.464 | 0.000 | 1.000 | 0.423 | 0.220 | 0.130 | 0.227 |
| 0.600 | 0.000 | 0.000 | 1.000 | 0.960 | 0.026 | 0.003 | 0.012 | 0.519 | 0.000 | 1.000 | 0.353 | 0.236 | 0.147 | 0.264 |
| 0.650 | 0.000 | 0.000 | 1.000 | 0.963 | 0.022 | 0.003 | 0.011 | 0.576 | 0.020 | 1.000 | 0.280 | 0.248 | 0.164 | 0.308 |
| 0.700 | 0.000 | 0.000 | 1.000 | 0.967 | 0.022 | 0.003 | 0.008 | 0.635 | 0.028 | 1.000 | 0.205 | 0.251 | 0.184 | 0.360 |
| 0.750 | 0.000 | 0.000 | 1.000 | 0.971 | 0.018 | 0.004 | 0.007 | 0.696 | 0.064 | 1.000 | 0.131 | 0.239 | 0.203 | 0.426 |
| 0.800 | 0.000 | 0.000 | 1.000 | 0.980 | 0.012 | 0.002 | 0.006 | 0.758 | 0.075 | 1.000 | 0.064 | 0.209 | 0.215 | 0.512 |
| 0.850 | 0.000 | 0.000 | 0.501 | 0.991 | 0.009 | 0.000 | 0.000 | 0.822 | 0.132 | 1.000 | 0.016 | 0.149 | 0.207 | 0.628 |
| 0.900 | 0.000 | 0.000 | 0.338 | 1.000 | 0.000 | 0.000 | 0.000 | 0.887 | 0.213 | 1.000 | 0.003 | 0.056 | 0.165 | 0.776 |
| 0.950 | 0.000 | 0.000 | 0.500 | 0.667 | 0.333 | 0.000 | 0.000 | 0.950 | 0.333 | 1.000 | 0.000 | 0.007 | 0.046 | 0.947 |

**Effects of thresholding on edge retention in the Glasser atlas.** For each absolute and relative threshold we show the proportion of edges that are consistently retained. As a measure of consistency, we use the number of participants in which the edge was retained at both timepoints divided by the ones in which it was retained at least once. For convenience, we then use the values defined in ^20^ to compute the ratio of edges having poor (ratio<0.40), fair (ratio=0.40-0.60), good (ratio=0.60-0.75) or excellent (ratio >0.75) consistency. For absolute thresholds, all edges below the value are set to 0, for relative ones (right) only the top percent corresponding to the threshold is retained.

Table S3

|  | Absolute | | | | | | | Relative | | | | | | |
| --- | --- | --- | --- | --- | --- | --- | --- | --- | --- | --- | --- | --- | --- | --- |
| Threshold | Median ratio | Min ratio | Max ratio | Poor edges ratio | Fair edges ratio | Good edges ratio | Excellent edges ratio | Median ratio | Min ratio | Max ratio | Poor edges ratio | Fair edges ratio | Good edges ratio | Excellent edges ratio |
| 0.05 | 0.473 | 0.023 | 1.000 | 0.385 | 0.280 | 0.154 | 0.182 | 0.065 | 0.000 | 1.000 | 0.918 | 0.050 | 0.015 | 0.017 |
| 0.10 | 0.307 | 0.000 | 1.000 | 0.646 | 0.198 | 0.083 | 0.074 | 0.075 | 0.000 | 1.000 | 0.895 | 0.060 | 0.023 | 0.022 |
| 0.15 | 0.223 | 0.000 | 1.000 | 0.792 | 0.133 | 0.041 | 0.034 | 0.100 | 0.000 | 1.000 | 0.858 | 0.079 | 0.031 | 0.032 |
| 0.20 | 0.183 | 0.000 | 1.000 | 0.876 | 0.084 | 0.023 | 0.018 | 0.132 | 0.000 | 1.000 | 0.812 | 0.101 | 0.042 | 0.045 |
| 0.25 | 0.151 | 0.000 | 1.000 | 0.921 | 0.055 | 0.012 | 0.011 | 0.168 | 0.000 | 1.000 | 0.765 | 0.121 | 0.054 | 0.059 |
| 0.30 | 0.118 | 0.000 | 1.000 | 0.944 | 0.039 | 0.009 | 0.008 | 0.209 | 0.000 | 1.000 | 0.716 | 0.140 | 0.068 | 0.076 |
| 0.35 | 0.070 | 0.000 | 1.000 | 0.955 | 0.032 | 0.006 | 0.008 | 0.253 | 0.000 | 1.000 | 0.668 | 0.156 | 0.081 | 0.095 |
| 0.40 | 0.000 | 0.000 | 1.000 | 0.960 | 0.028 | 0.004 | 0.008 | 0.299 | 0.000 | 1.000 | 0.618 | 0.171 | 0.096 | 0.116 |
| 0.45 | 0.000 | 0.000 | 1.000 | 0.962 | 0.025 | 0.004 | 0.009 | 0.346 | 0.000 | 1.000 | 0.564 | 0.187 | 0.109 | 0.140 |
| 0.50 | 0.000 | 0.000 | 1.000 | 0.961 | 0.024 | 0.003 | 0.011 | 0.395 | 0.009 | 1.000 | 0.505 | 0.205 | 0.123 | 0.167 |
| 0.55 | 0.000 | 0.000 | 1.000 | 0.962 | 0.023 | 0.004 | 0.011 | 0.446 | 0.037 | 1.000 | 0.442 | 0.223 | 0.135 | 0.200 |
| 0.60 | 0.000 | 0.000 | 1.000 | 0.961 | 0.024 | 0.002 | 0.012 | 0.498 | 0.047 | 1.000 | 0.372 | 0.243 | 0.150 | 0.235 |
| 0.65 | 0.000 | 0.000 | 1.000 | 0.959 | 0.028 | 0.002 | 0.011 | 0.553 | 0.085 | 1.000 | 0.295 | 0.262 | 0.165 | 0.278 |
| 0.70 | 0.000 | 0.000 | 1.000 | 0.960 | 0.019 | 0.003 | 0.018 | 0.610 | 0.108 | 1.000 | 0.205 | 0.281 | 0.186 | 0.328 |
| 0.75 | 0.000 | 0.000 | 1.000 | 0.980 | 0.013 | 0.004 | 0.004 | 0.670 | 0.141 | 1.000 | 0.116 | 0.283 | 0.211 | 0.390 |
| 0.80 | 0.000 | 0.000 | 1.000 | 0.954 | 0.029 | 0.004 | 0.013 | 0.731 | 0.179 | 1.000 | 0.042 | 0.246 | 0.240 | 0.472 |
| 0.85 | 0.000 | 0.000 | 1.000 | 0.968 | 0.021 | 0.000 | 0.011 | 0.797 | 0.240 | 1.000 | 0.004 | 0.150 | 0.256 | 0.590 |
| 0.90 | 0.000 | 0.000 | 0.356 | 1.000 | 0.000 | 0.000 | 0.000 | 0.866 | 0.324 | 1.000 | 0.000 | 0.036 | 0.190 | 0.773 |
| 0.95 | 1.000 | 1.000 | 1.000 | 0.000 | 0.000 | 0.000 | 1.000 | 0.937 | 0.440 | 1.000 | 0.000 | 0.002 | 0.035 | 0.963 |

**Effects of thresholding on edge retention in the Gordon atlas.** For each absolute and relative threshold we show the proportion of edges that are consistently retained. As a measure of consistency, we use the number of participants in which the edge was retained at both timepoints divided by the ones in which it was retained at least once. For convenience, we then use the values defined in ^20^ to compute the ratio of edges having poor (ratio<0.40), fair (ratio=0.40-0.60), good (ratio=0.60-0.75) or excellent (ratio >0.75) consistency. For absolute thresholds, all edges below the value are set to 0, for relative ones (right) only the top percent corresponding to the threshold is retained.

Table S4

|  | Absolute | | | | | | | Relative | | | | | | |
| --- | --- | --- | --- | --- | --- | --- | --- | --- | --- | --- | --- | --- | --- | --- |
| Threshold | Median ICC | Min ICC | Max ICC | Poor edges ratio | Fair edges ratio | Good edges ratio | Excellent edges ratio | Median ICC | Min ICC | Max ICC | Poor edges ratio | Fair edges ratio | Good edges ratio | Excellent edges ratio |
| 0.050 | 0.391 | -0.320 | 0.808 | 0.531 | 0.429 | 0.040 | 0.000 | 0.525 | -1.000 | 1.000 | 0.247 | 0.447 | 0.182 | 0.124 |
| 0.100 | 0.367 | -0.956 | 0.997 | 0.609 | 0.359 | 0.031 | 0.001 | 0.525 | -1.000 | 1.000 | 0.219 | 0.495 | 0.176 | 0.111 |
| 0.150 | 0.346 | -1.000 | 0.999 | 0.669 | 0.299 | 0.027 | 0.005 | 0.523 | -1.000 | 1.000 | 0.199 | 0.540 | 0.167 | 0.094 |
| 0.200 | 0.328 | -1.000 | 1.000 | 0.712 | 0.256 | 0.023 | 0.009 | 0.520 | -1.000 | 1.000 | 0.178 | 0.587 | 0.160 | 0.076 |
| 0.250 | 0.311 | -1.000 | 1.000 | 0.747 | 0.219 | 0.022 | 0.012 | 0.514 | -1.000 | 1.000 | 0.169 | 0.626 | 0.143 | 0.063 |
| 0.300 | 0.293 | -1.000 | 1.000 | 0.767 | 0.195 | 0.023 | 0.015 | 0.508 | -1.000 | 1.000 | 0.163 | 0.655 | 0.127 | 0.055 |
| 0.350 | 0.275 | -1.000 | 1.000 | 0.774 | 0.178 | 0.027 | 0.020 | 0.503 | -1.000 | 1.000 | 0.158 | 0.681 | 0.117 | 0.044 |
| 0.400 | 0.260 | -1.000 | 1.000 | 0.768 | 0.170 | 0.034 | 0.028 | 0.497 | -1.000 | 1.000 | 0.162 | 0.699 | 0.103 | 0.036 |
| 0.450 | 0.247 | -1.000 | 1.000 | 0.764 | 0.155 | 0.039 | 0.043 | 0.490 | -1.000 | 1.000 | 0.168 | 0.714 | 0.090 | 0.027 |
| 0.500 | 0.236 | -1.000 | 1.000 | 0.752 | 0.150 | 0.045 | 0.052 | 0.484 | -1.000 | 1.000 | 0.172 | 0.729 | 0.080 | 0.019 |
| 0.550 | 0.235 | -1.000 | 1.000 | 0.743 | 0.145 | 0.048 | 0.064 | 0.478 | -1.000 | 0.997 | 0.176 | 0.744 | 0.070 | 0.010 |
| 0.600 | 0.228 | -1.000 | 1.000 | 0.738 | 0.142 | 0.047 | 0.074 | 0.471 | -0.931 | 0.999 | 0.184 | 0.751 | 0.060 | 0.004 |
| 0.650 | 0.224 | -1.000 | 1.000 | 0.734 | 0.137 | 0.050 | 0.080 | 0.465 | -0.433 | 0.970 | 0.197 | 0.750 | 0.051 | 0.002 |
| 0.700 | 0.222 | -1.000 | 1.000 | 0.730 | 0.142 | 0.047 | 0.080 | 0.459 | -0.269 | 0.827 | 0.214 | 0.737 | 0.049 | 0.001 |
| 0.750 | 0.214 | -1.000 | 1.000 | 0.720 | 0.139 | 0.053 | 0.088 | 0.454 | -0.155 | 0.807 | 0.240 | 0.712 | 0.048 | 0.001 |
| 0.800 | 0.229 | -0.996 | 1.000 | 0.731 | 0.131 | 0.047 | 0.090 | 0.450 | -0.101 | 0.808 | 0.270 | 0.679 | 0.051 | 0.001 |
| 0.850 | 0.239 | -1.000 | 0.994 | 0.661 | 0.115 | 0.067 | 0.158 | 0.447 | -0.048 | 0.808 | 0.297 | 0.648 | 0.054 | 0.001 |
| 0.900 | 0.188 | -0.999 | 0.848 | 0.743 | 0.171 | 0.029 | 0.057 | 0.447 | -0.046 | 0.810 | 0.314 | 0.626 | 0.060 | 0.001 |
| 0.950 | 0.579 | 0.378 | 0.780 | 0.500 | 0.000 | 0.000 | 0.500 | 0.452 | -0.036 | 0.808 | 0.311 | 0.622 | 0.065 | 0.001 |

**Effects of thresholding on ICC in the Brainnetome atlas.** for each absolute and relative threshold we show the proportion of edges having poor (ICC<0.40), fair (ICC=0.40-0.60), good (ICC=0.60-0.75) or excellent (ICC>0.75) reliability. In this calculation, only subjects for which the edge was retained in both sessions were considered. For absolute thresholds (left) all edges below the value are set to 0, for relative ones (right) only the top percent corresponding to the threshold is retained. Abbreviations: ICC=intraclass correlation coefficient.

Table S5

|  | Absolute | | | | | | | Relative | | | | | | |
| --- | --- | --- | --- | --- | --- | --- | --- | --- | --- | --- | --- | --- | --- | --- |
| Threshold | Median ICC | Min ICC | Max ICC | Poor edges ratio | Fair edges ratio | Good edges ratio | Excellent edges ratio | Median ICC | Min ICC | Max ICC | Poor edges ratio | Fair edges ratio | Good edges ratio | Excellent edges ratio |
| 0.050 | 0.390 | -0.568 | 0.823 | 0.536 | 0.417 | 0.046 | 0.001 | 0.588 | -1.000 | 1.000 | 0.174 | 0.364 | 0.305 | 0.158 |
| 0.100 | 0.369 | -1.000 | 1.000 | 0.601 | 0.355 | 0.039 | 0.005 | 0.573 | -1.000 | 1.000 | 0.167 | 0.422 | 0.272 | 0.140 |
| 0.150 | 0.342 | -1.000 | 1.000 | 0.656 | 0.293 | 0.038 | 0.013 | 0.561 | -1.000 | 1.000 | 0.157 | 0.469 | 0.250 | 0.123 |
| 0.200 | 0.312 | -1.000 | 1.000 | 0.683 | 0.247 | 0.043 | 0.026 | 0.554 | -1.000 | 1.000 | 0.148 | 0.506 | 0.234 | 0.112 |
| 0.250 | 0.287 | -1.000 | 1.000 | 0.692 | 0.219 | 0.048 | 0.041 | 0.547 | -1.000 | 1.000 | 0.140 | 0.545 | 0.219 | 0.097 |
| 0.300 | 0.265 | -1.000 | 1.000 | 0.695 | 0.198 | 0.052 | 0.055 | 0.540 | -1.000 | 1.000 | 0.131 | 0.581 | 0.205 | 0.082 |
| 0.350 | 0.253 | -1.000 | 1.000 | 0.692 | 0.190 | 0.052 | 0.066 | 0.533 | -1.000 | 1.000 | 0.123 | 0.622 | 0.191 | 0.064 |
| 0.400 | 0.251 | -1.000 | 1.000 | 0.684 | 0.185 | 0.058 | 0.073 | 0.526 | -1.000 | 1.000 | 0.120 | 0.657 | 0.176 | 0.047 |
| 0.450 | 0.252 | -1.000 | 1.000 | 0.674 | 0.181 | 0.059 | 0.085 | 0.518 | -0.999 | 1.000 | 0.115 | 0.701 | 0.157 | 0.028 |
| 0.500 | 0.244 | -1.000 | 1.000 | 0.674 | 0.174 | 0.061 | 0.090 | 0.511 | -0.999 | 1.000 | 0.112 | 0.739 | 0.135 | 0.014 |
| 0.550 | 0.255 | -1.000 | 1.000 | 0.658 | 0.166 | 0.067 | 0.109 | 0.503 | -0.992 | 0.998 | 0.113 | 0.768 | 0.112 | 0.007 |
| 0.600 | 0.263 | -1.000 | 1.000 | 0.656 | 0.179 | 0.067 | 0.097 | 0.496 | -0.921 | 0.998 | 0.117 | 0.787 | 0.092 | 0.004 |
| 0.650 | 0.267 | -0.999 | 1.000 | 0.666 | 0.162 | 0.056 | 0.115 | 0.488 | -0.362 | 0.932 | 0.126 | 0.795 | 0.077 | 0.002 |
| 0.700 | 0.271 | -1.000 | 0.994 | 0.680 | 0.168 | 0.063 | 0.090 | 0.480 | -0.190 | 0.869 | 0.141 | 0.790 | 0.068 | 0.001 |
| 0.750 | 0.252 | -0.996 | 0.992 | 0.686 | 0.168 | 0.044 | 0.102 | 0.472 | -0.095 | 0.896 | 0.162 | 0.772 | 0.065 | 0.001 |
| 0.800 | 0.229 | -0.782 | 0.977 | 0.765 | 0.118 | 0.020 | 0.098 | 0.465 | -0.028 | 0.836 | 0.190 | 0.745 | 0.064 | 0.001 |
| 0.850 | 0.434 | -0.970 | 0.868 | 0.421 | 0.368 | 0.105 | 0.105 | 0.458 | -0.067 | 0.823 | 0.226 | 0.707 | 0.066 | 0.001 |
| 0.900 | 0.416 | -0.295 | 0.764 | 0.500 | 0.250 | 0.000 | 0.250 | 0.453 | -0.051 | 0.823 | 0.264 | 0.667 | 0.068 | 0.001 |
| 0.950 | - | - | - | - | - | - | - | 0.452 | -0.034 | 0.824 | 0.295 | 0.632 | 0.071 | 0.001 |

**Effects of thresholding on ICC in the Glasser atlas.** for each absolute and relative threshold we show the proportion of edges having poor (ICC<0.40), fair (ICC=0.40-0.60), good (ICC=0.60-0.75) or excellent (ICC>0.75) reliability. In this calculation, only subjects for which the edge was retained in both sessions were considered. For absolute thresholds (left) all edges below the value are set to 0, for relative ones (right) only the top percent corresponding to the threshold is retained. A dash indicates that ICC could not be computed for that threshold (not enough remaining consistent edges). Abbreviations: ICC=intraclass correlation coefficient.

Table S6

|  | Absolute | | | | | | | Relative | | | | | | |
| --- | --- | --- | --- | --- | --- | --- | --- | --- | --- | --- | --- | --- | --- | --- |
| Threshold | Median ICC | Min ICC | Max ICC | Poor edges ratio | Fair edges ratio | Good edges ratio | Excellent edges ratio | Median ICC | Min ICC | Max ICC | Poor edges ratio | Fair edges ratio | Good edges ratio | Excellent edges ratio |
| 0.050 | 0.356 | -0.794 | 0.888 | 0.643 | 0.327 | 0.030 | 0.000 | 0.571 | -1.000 | 1.000 | 0.200 | 0.383 | 0.264 | 0.153 |
| 0.100 | 0.335 | -1.000 | 1.000 | 0.678 | 0.282 | 0.031 | 0.009 | 0.553 | -1.000 | 1.000 | 0.190 | 0.450 | 0.235 | 0.125 |
| 0.150 | 0.307 | -1.000 | 1.000 | 0.707 | 0.237 | 0.035 | 0.021 | 0.541 | -1.000 | 1.000 | 0.182 | 0.498 | 0.207 | 0.113 |
| 0.200 | 0.283 | -1.000 | 1.000 | 0.714 | 0.208 | 0.041 | 0.036 | 0.531 | -1.000 | 1.000 | 0.177 | 0.535 | 0.191 | 0.097 |
| 0.250 | 0.260 | -1.000 | 1.000 | 0.712 | 0.192 | 0.047 | 0.049 | 0.522 | -1.000 | 1.000 | 0.176 | 0.563 | 0.175 | 0.086 |
| 0.300 | 0.245 | -1.000 | 1.000 | 0.703 | 0.179 | 0.054 | 0.063 | 0.511 | -1.000 | 1.000 | 0.186 | 0.586 | 0.160 | 0.069 |
| 0.350 | 0.230 | -1.000 | 1.000 | 0.697 | 0.171 | 0.058 | 0.074 | 0.501 | -1.000 | 1.000 | 0.193 | 0.620 | 0.142 | 0.045 |
| 0.400 | 0.236 | -1.000 | 1.000 | 0.685 | 0.169 | 0.062 | 0.085 | 0.491 | -1.000 | 1.000 | 0.198 | 0.656 | 0.121 | 0.025 |
| 0.450 | 0.230 | -1.000 | 1.000 | 0.681 | 0.160 | 0.063 | 0.096 | 0.481 | -0.993 | 1.000 | 0.205 | 0.688 | 0.096 | 0.011 |
| 0.500 | 0.258 | -1.000 | 1.000 | 0.667 | 0.159 | 0.069 | 0.104 | 0.472 | -0.859 | 0.992 | 0.219 | 0.702 | 0.075 | 0.004 |
| 0.550 | 0.270 | -1.000 | 1.000 | 0.644 | 0.178 | 0.064 | 0.114 | 0.463 | -0.482 | 0.962 | 0.239 | 0.703 | 0.057 | 0.001 |
| 0.600 | 0.281 | -1.000 | 1.000 | 0.661 | 0.156 | 0.073 | 0.109 | 0.453 | -0.404 | 0.863 | 0.269 | 0.683 | 0.048 | 0.000 |
| 0.650 | 0.246 | -0.992 | 0.996 | 0.690 | 0.144 | 0.065 | 0.101 | 0.443 | -0.279 | 0.790 | 0.304 | 0.651 | 0.044 | 0.000 |
| 0.700 | 0.221 | -0.997 | 0.999 | 0.663 | 0.134 | 0.081 | 0.122 | 0.434 | -0.205 | 0.791 | 0.347 | 0.610 | 0.043 | 0.000 |
| 0.750 | 0.162 | -1.000 | 0.960 | 0.701 | 0.130 | 0.065 | 0.104 | 0.425 | -0.173 | 0.795 | 0.390 | 0.565 | 0.044 | 0.000 |
| 0.800 | 0.169 | -0.987 | 0.960 | 0.640 | 0.160 | 0.080 | 0.120 | 0.417 | -0.119 | 0.798 | 0.432 | 0.522 | 0.045 | 0.001 |
| 0.850 | 0.249 | 0.123 | 0.475 | 0.750 | 0.250 | 0.000 | 0.000 | 0.410 | -0.122 | 0.801 | 0.463 | 0.489 | 0.047 | 0.001 |
| 0.900 | 0.062 | -0.295 | 0.420 | 0.500 | 0.500 | 0.000 | 0.000 | 0.405 | -0.067 | 0.804 | 0.482 | 0.468 | 0.049 | 0.001 |
| 0.950 | - | - | - | - | - | - | - | 0.404 | -0.062 | 0.806 | 0.487 | 0.461 | 0.051 | 0.001 |

**Effects of thresholding on ICC in the Gordon atlas.** for each absolute and relative threshold we show the proportion of edges having poor (ICC<0.40), fair (ICC=0.40-0.60), good (ICC=0.60-0.75) or excellent (ICC>0.75) reliability. In this calculation, only subjects for which the edge was retained in both sessions were considered. For absolute thresholds (left) all edges below the value are set to 0, for relative ones (right) only the top percent corresponding to the threshold is retained. A dash indicates that ICC could not be computed for that threshold (not enough remaining consistent edges). Abbreviations: ICC=intraclass correlation coefficient.

Table S7

|  |  | Median ICC | Min ICC | Max ICC | Poor edges ratio | Fair edges ratio | Good edges ratio | Excellent edges ratio |
| --- | --- | --- | --- | --- | --- | --- | --- | --- |
| BNT | GSR- | 0.474 | -0.445 | 0.838 | 0.330 | 0.502 | 0.163 | 0.005 |
|  | GSR+ | 0.382 | -0.447 | 0.896 | 0.534 | 0.313 | 0.125 | 0.029 |
| Glasser | GSR- | 0.457 | -0.297 | 0.870 | 0.355 | 0.511 | 0.131 | 0.003 |
|  | GSR+ | 0.344 | -0.297 | 0.900 | 0.610 | 0.297 | 0.085 | 0.009 |
| Gordon | GSR- | 0.412 | -0.350 | 0.825 | 0.473 | 0.430 | 0.094 | 0.002 |
|  | GSR+ | 0.305 | -0.345 | 0.881 | 0.674 | 0.245 | 0.073 | 0.008 |

**Reliability of edges in the functional connectome in unrelated participants.** We show the median, minimum and maximum ICC of functional connectomes computed using three different atlases, with or without global signal regression. We also show the proportion of edges having poor (ICC<0.40), fair (ICC=0.40-0.60), good (ICC=0.60-0.75) or excellent (ICC>0.75) reliability, defined in accordance to ^20^. Abbreviations: ICC=intraclass correlation coefficient, GSR-=no global signal regression, GSR+=global signal regression.

Table S8

|  | ICC | | Lower CI | | Upper CI | |
| --- | --- | --- | --- | --- | --- | --- |
|  | GSR- | GSR+ | GSR- | GSR+ | GSR- | GSR+ |
| DMN | 0.514 | 0.540 | 0.336 | 0.676 | 0.656 | 0.368 |
| PAO | 0.691 | 0.759 | 0.559 | 0.837 | 0.788 | 0.651 |
| FP | 0.423 | 0.532 | 0.229 | 0.670 | 0.585 | 0.359 |
| SAL | 0.583 | 0.528 | 0.421 | 0.667 | 0.709 | 0.354 |
| COP | 0.636 | 0.505 | 0.488 | 0.649 | 0.748 | 0.326 |
| MEP | 0.645 | 0.718 | 0.499 | 0.808 | 0.755 | 0.595 |
| DAN | 0.587 | 0.492 | 0.426 | 0.639 | 0.712 | 0.310 |
| VAN | 0.593 | 0.652 | 0.434 | 0.761 | 0.717 | 0.509 |
| VIS | 0.697 | 0.674 | 0.568 | 0.776 | 0.793 | 0.536 |
| SMH | 0.741 | 0.596 | 0.626 | 0.719 | 0.825 | 0.438 |
| SMM | 0.743 | 0.452 | 0.628 | 0.608 | 0.826 | 0.263 |
| AUD | 0.755 | 0.572 | 0.645 | 0.700 | 0.834 | 0.407 |

**Reliability of known resting state networks in unrelated participants.** We show the ICC and confidence intervals for the average connectivity within known resting state networks defined in accordance to ^28^. Abbreviations: ICC=intraclass correlation coefficient, GSR-=no global signal regression, GSR+=global signal regression, CI=confidence interval, DMN=default mode network, PAO=parieto-occipital, FP=fronto-parietal, SAL=salience, COP=cingulo-opercular, MEP=medial parietal, DAN=dorsal attention network, VAN=ventral attention network, VIS=visual, SMH=supplementary motor (hand), SMM=supplementary motor (mouth), AUD=auditory.

Table S9

|  | Absolute | | | | | | | Relative | | | | | | |
| --- | --- | --- | --- | --- | --- | --- | --- | --- | --- | --- | --- | --- | --- | --- |
| Threshold | Median ratio | Min ratio | Max ratio | Poor edges ratio | Fair edges ratio | Good edges ratio | Excellent edges ratio | Median ratio | Min ratio | Max ratio | Poor edges ratio | Fair edges ratio | Good edges ratio | Excellent edges ratio |
| 0.050 | 0.750 | 0.000 | 1.000 | 0.107 | 0.187 | 0.202 | 0.504 | 0.143 | 0.000 | 1.000 | 0.773 | 0.141 | 0.043 | 0.043 |
| 0.100 | 0.629 | 0.000 | 1.000 | 0.228 | 0.232 | 0.192 | 0.348 | 0.192 | 0.000 | 1.000 | 0.743 | 0.153 | 0.054 | 0.050 |
| 0.150 | 0.519 | 0.000 | 1.000 | 0.345 | 0.254 | 0.168 | 0.234 | 0.206 | 0.000 | 1.000 | 0.717 | 0.157 | 0.061 | 0.064 |
| 0.200 | 0.431 | 0.000 | 1.000 | 0.452 | 0.251 | 0.142 | 0.155 | 0.231 | 0.000 | 1.000 | 0.689 | 0.159 | 0.073 | 0.079 |
| 0.250 | 0.370 | 0.000 | 1.000 | 0.538 | 0.241 | 0.117 | 0.104 | 0.250 | 0.000 | 1.000 | 0.655 | 0.166 | 0.082 | 0.098 |
| 0.300 | 0.327 | 0.000 | 1.000 | 0.614 | 0.223 | 0.090 | 0.073 | 0.286 | 0.000 | 1.000 | 0.618 | 0.174 | 0.088 | 0.120 |
| 0.350 | 0.286 | 0.000 | 1.000 | 0.675 | 0.200 | 0.073 | 0.052 | 0.326 | 0.000 | 1.000 | 0.578 | 0.179 | 0.099 | 0.144 |
| 0.400 | 0.250 | 0.000 | 1.000 | 0.720 | 0.183 | 0.058 | 0.039 | 0.364 | 0.000 | 1.000 | 0.534 | 0.183 | 0.109 | 0.173 |
| 0.450 | 0.222 | 0.000 | 1.000 | 0.748 | 0.176 | 0.044 | 0.033 | 0.414 | 0.000 | 1.000 | 0.483 | 0.191 | 0.120 | 0.206 |
| 0.500 | 0.200 | 0.000 | 1.000 | 0.774 | 0.162 | 0.035 | 0.029 | 0.467 | 0.000 | 1.000 | 0.435 | 0.191 | 0.131 | 0.243 |
| 0.550 | 0.143 | 0.000 | 1.000 | 0.797 | 0.148 | 0.026 | 0.030 | 0.521 | 0.000 | 1.000 | 0.380 | 0.194 | 0.142 | 0.284 |
| 0.600 | 0.059 | 0.000 | 1.000 | 0.805 | 0.141 | 0.025 | 0.028 | 0.578 | 0.000 | 1.000 | 0.326 | 0.193 | 0.152 | 0.328 |
| 0.650 | 0.000 | 0.000 | 1.000 | 0.816 | 0.125 | 0.026 | 0.033 | 0.634 | 0.000 | 1.000 | 0.271 | 0.192 | 0.158 | 0.379 |
| 0.700 | 0.000 | 0.000 | 1.000 | 0.834 | 0.113 | 0.022 | 0.032 | 0.692 | 0.000 | 1.000 | 0.215 | 0.184 | 0.166 | 0.435 |
| 0.750 | 0.000 | 0.000 | 1.000 | 0.818 | 0.123 | 0.030 | 0.029 | 0.747 | 0.000 | 1.000 | 0.156 | 0.177 | 0.170 | 0.497 |
| 0.800 | 0.000 | 0.000 | 1.000 | 0.837 | 0.095 | 0.024 | 0.043 | 0.800 | 0.050 | 1.000 | 0.095 | 0.162 | 0.171 | 0.572 |
| 0.850 | 0.000 | 0.000 | 1.000 | 0.812 | 0.136 | 0.024 | 0.028 | 0.852 | 0.077 | 1.000 | 0.035 | 0.136 | 0.166 | 0.663 |
| 0.900 | 0.000 | 0.000 | 1.000 | 0.867 | 0.117 | 0.000 | 0.017 | 0.904 | 0.100 | 1.000 | 0.004 | 0.070 | 0.144 | 0.782 |
| 0.950 | 0.167 | 0.000 | 0.500 | 0.750 | 0.250 | 0.000 | 0.000 | 0.952 | 0.200 | 1.000 | 0.000 | 0.007 | 0.054 | 0.940 |

**Effects of thresholding on edge retention in the Brainnetome atlas in unrelated participants.** For each absolute and relative threshold we show the proportion of edges that are consistently retained. As a measure of consistency, we use the number of participants in which the edge was retained at both timepoints divided by the ones in which it was retained at least once. For convenience, we then use the values defined in ^20^ to compute the ratio of edges having poor (ratio<0.40), fair (ratio=0.40-0.60), good (ratio=0.60-0.75) or excellent (ratio >0.75) consistency. For absolute thresholds, all edges below the value are set to 0, for relative ones (right) only the top percent corresponding to the threshold is retained.

Table S10

|  | Absolute | | | | | | | Relative | | | | | | |
| --- | --- | --- | --- | --- | --- | --- | --- | --- | --- | --- | --- | --- | --- | --- |
| Threshold | Median ratio | Min ratio | Max ratio | Poor edges ratio | Fair edges ratio | Good edges ratio | Excellent edges ratio | Median ratio | Min ratio | Max ratio | Poor edges ratio | Fair edges ratio | Good edges ratio | Excellent edges ratio |
| 0.050 | 0.578 | 0.000 | 1.000 | 0.220 | 0.310 | 0.192 | 0.278 | 0.125 | 0.000 | 1.000 | 0.792 | 0.129 | 0.035 | 0.043 |
| 0.100 | 0.412 | 0.000 | 1.000 | 0.474 | 0.273 | 0.119 | 0.133 | 0.143 | 0.000 | 1.000 | 0.786 | 0.122 | 0.045 | 0.048 |
| 0.150 | 0.324 | 0.000 | 1.000 | 0.627 | 0.225 | 0.077 | 0.071 | 0.154 | 0.000 | 1.000 | 0.776 | 0.119 | 0.048 | 0.057 |
| 0.200 | 0.250 | 0.000 | 1.000 | 0.721 | 0.188 | 0.051 | 0.040 | 0.172 | 0.000 | 1.000 | 0.760 | 0.117 | 0.055 | 0.068 |
| 0.250 | 0.200 | 0.000 | 1.000 | 0.786 | 0.152 | 0.035 | 0.026 | 0.200 | 0.000 | 1.000 | 0.730 | 0.125 | 0.062 | 0.083 |
| 0.300 | 0.133 | 0.000 | 1.000 | 0.824 | 0.130 | 0.026 | 0.021 | 0.235 | 0.000 | 1.000 | 0.691 | 0.140 | 0.068 | 0.101 |
| 0.350 | 0.000 | 0.000 | 1.000 | 0.849 | 0.110 | 0.020 | 0.021 | 0.273 | 0.000 | 1.000 | 0.648 | 0.153 | 0.079 | 0.120 |
| 0.400 | 0.000 | 0.000 | 1.000 | 0.870 | 0.092 | 0.016 | 0.022 | 0.316 | 0.000 | 1.000 | 0.597 | 0.170 | 0.091 | 0.142 |
| 0.450 | 0.000 | 0.000 | 1.000 | 0.878 | 0.085 | 0.013 | 0.024 | 0.361 | 0.000 | 1.000 | 0.542 | 0.187 | 0.103 | 0.168 |
| 0.500 | 0.000 | 0.000 | 1.000 | 0.882 | 0.077 | 0.013 | 0.027 | 0.411 | 0.000 | 1.000 | 0.484 | 0.204 | 0.116 | 0.196 |
| 0.550 | 0.000 | 0.000 | 1.000 | 0.874 | 0.082 | 0.013 | 0.031 | 0.463 | 0.000 | 1.000 | 0.422 | 0.219 | 0.130 | 0.229 |
| 0.600 | 0.000 | 0.000 | 1.000 | 0.869 | 0.078 | 0.014 | 0.038 | 0.517 | 0.000 | 1.000 | 0.353 | 0.235 | 0.145 | 0.267 |
| 0.650 | 0.000 | 0.000 | 1.000 | 0.858 | 0.076 | 0.018 | 0.047 | 0.575 | 0.000 | 1.000 | 0.283 | 0.244 | 0.163 | 0.310 |
| 0.700 | 0.000 | 0.000 | 1.000 | 0.837 | 0.092 | 0.023 | 0.048 | 0.635 | 0.000 | 1.000 | 0.207 | 0.248 | 0.182 | 0.363 |
| 0.750 | 0.000 | 0.000 | 1.000 | 0.780 | 0.154 | 0.017 | 0.050 | 0.696 | 0.000 | 1.000 | 0.135 | 0.236 | 0.199 | 0.430 |
| 0.800 | 0.200 | 0.000 | 0.678 | 0.881 | 0.071 | 0.048 | 0.000 | 0.759 | 0.000 | 1.000 | 0.072 | 0.202 | 0.209 | 0.517 |
| 0.850 | 0.000 | 0.000 | 0.571 | 0.882 | 0.118 | 0.000 | 0.000 | 0.825 | 0.057 | 1.000 | 0.025 | 0.143 | 0.202 | 0.629 |
| 0.900 | 0.083 | 0.000 | 0.500 | 0.833 | 0.167 | 0.000 | 0.000 | 0.890 | 0.152 | 1.000 | 0.005 | 0.063 | 0.158 | 0.775 |
| 0.950 | - | - | - | - | - | - | - | 0.952 | 0.266 | 1.000 | 0.000 | 0.009 | 0.052 | 0.938 |

**Effects of thresholding on edge retention in the Glasser atlas in unrelated participants.** For each absolute and relative threshold we show the proportion of edges that are consistently retained. As a measure of consistency, we use the number of participants in which the edge was retained at both timepoints divided by the ones in which it was retained at least once. For convenience, we then use the values defined in ^20^ to compute the ratio of edges having poor (ratio<0.40), fair (ratio=0.40-0.60), good (ratio=0.60-0.75) or excellent (ratio >0.75) consistency. For absolute thresholds, all edges below the value are set to 0, for relative ones (right) only the top percent corresponding to the threshold is retained. A dash indicates that consistency could not be computed for that threshold (not enough remaining consistent edges).

Table S11

|  | Absolute | | | | | | | Relative | | | | | | |
| --- | --- | --- | --- | --- | --- | --- | --- | --- | --- | --- | --- | --- | --- | --- |
| Threshold | Median ratio | Min ratio | Max ratio | Poor edges ratio | Fair edges ratio | Good edges ratio | Excellent edges ratio | Median ratio | Min ratio | Max ratio | Poor edges ratio | Fair edges ratio | Good edges ratio | Excellent edges ratio |
| 0.050 | 0.479 | 0.000 | 1.000 | 0.370 | 0.291 | 0.154 | 0.184 | 0.105 | 0.000 | 1.000 | 0.820 | 0.113 | 0.028 | 0.038 |
| 0.100 | 0.333 | 0.000 | 1.000 | 0.608 | 0.225 | 0.089 | 0.077 | 0.118 | 0.000 | 1.000 | 0.819 | 0.108 | 0.036 | 0.037 |
| 0.150 | 0.267 | 0.000 | 1.000 | 0.715 | 0.193 | 0.052 | 0.040 | 0.133 | 0.000 | 1.000 | 0.806 | 0.112 | 0.040 | 0.043 |
| 0.200 | 0.200 | 0.000 | 1.000 | 0.784 | 0.160 | 0.031 | 0.025 | 0.160 | 0.000 | 1.000 | 0.781 | 0.120 | 0.046 | 0.053 |
| 0.250 | 0.143 | 0.000 | 1.000 | 0.832 | 0.126 | 0.021 | 0.021 | 0.194 | 0.000 | 1.000 | 0.747 | 0.132 | 0.057 | 0.064 |
| 0.300 | 0.000 | 0.000 | 1.000 | 0.861 | 0.103 | 0.016 | 0.019 | 0.227 | 0.000 | 1.000 | 0.706 | 0.145 | 0.068 | 0.080 |
| 0.350 | 0.000 | 0.000 | 1.000 | 0.877 | 0.089 | 0.013 | 0.021 | 0.263 | 0.000 | 1.000 | 0.660 | 0.160 | 0.081 | 0.099 |
| 0.400 | 0.000 | 0.000 | 1.000 | 0.887 | 0.078 | 0.012 | 0.024 | 0.306 | 0.000 | 1.000 | 0.611 | 0.174 | 0.095 | 0.119 |
| 0.450 | 0.000 | 0.000 | 1.000 | 0.893 | 0.070 | 0.011 | 0.026 | 0.351 | 0.000 | 1.000 | 0.558 | 0.191 | 0.108 | 0.143 |
| 0.500 | 0.000 | 0.000 | 1.000 | 0.893 | 0.069 | 0.010 | 0.027 | 0.400 | 0.000 | 1.000 | 0.499 | 0.209 | 0.121 | 0.171 |
| 0.550 | 0.000 | 0.000 | 1.000 | 0.890 | 0.068 | 0.010 | 0.032 | 0.448 | 0.000 | 1.000 | 0.437 | 0.227 | 0.135 | 0.202 |
| 0.600 | 0.000 | 0.000 | 1.000 | 0.883 | 0.072 | 0.010 | 0.035 | 0.500 | 0.000 | 1.000 | 0.366 | 0.247 | 0.150 | 0.237 |
| 0.650 | 0.000 | 0.000 | 1.000 | 0.860 | 0.094 | 0.010 | 0.037 | 0.554 | 0.000 | 1.000 | 0.288 | 0.266 | 0.166 | 0.280 |
| 0.700 | 0.000 | 0.000 | 1.000 | 0.852 | 0.085 | 0.012 | 0.051 | 0.611 | 0.000 | 1.000 | 0.206 | 0.278 | 0.187 | 0.329 |
| 0.750 | 0.000 | 0.000 | 1.000 | 0.844 | 0.078 | 0.000 | 0.078 | 0.671 | 0.051 | 1.000 | 0.124 | 0.271 | 0.213 | 0.392 |
| 0.800 | 0.000 | 0.000 | 1.000 | 0.778 | 0.067 | 0.044 | 0.111 | 0.734 | 0.111 | 1.000 | 0.053 | 0.232 | 0.239 | 0.476 |
| 0.850 | 0.000 | 0.000 | 0.467 | 0.818 | 0.182 | 0.000 | 0.000 | 0.800 | 0.160 | 1.000 | 0.011 | 0.150 | 0.241 | 0.598 |
| 0.900 | 0.000 | 0.000 | 0.231 | 1.000 | 0.000 | 0.000 | 0.000 | 0.867 | 0.227 | 1.000 | 0.001 | 0.048 | 0.179 | 0.772 |
| 0.950 | - | - | - | - | - | - | - | 0.940 | 0.352 | 1.000 | 0.000 | 0.004 | 0.045 | 0.951 |

**Effects of thresholding on edge retention in the Gordon atlas in unrelated participants.** For each absolute and relative threshold we show the proportion of edges that are consistently retained. As a measure of consistency, we use the number of participants in which the edge was retained at both timepoints divided by the ones in which it was retained at least once. For convenience, we then use the values defined in ^20^ to compute the ratio of edges having poor (ratio<0.40), fair (ratio=0.40-0.60), good (ratio=0.60-0.75) or excellent (ratio >0.75) consistency. For absolute thresholds, all edges below the value are set to 0, for relative ones (right) only the top percent corresponding to the threshold is retained. A dash indicates that consistency could not be computed for that threshold (not enough remaining consistent edges).

Table S12

|  | Absolute | | | | | | | Relative | | | | | | |
| --- | --- | --- | --- | --- | --- | --- | --- | --- | --- | --- | --- | --- | --- | --- |
| Threshold | Median ICC | Min ICC | Max ICC | Poor edges ratio | Fair edges ratio | Good edges ratio | Excellent edges ratio | Median ICC | Min ICC | Max ICC | Poor edges ratio | Fair edges ratio | Good edges ratio | Excellent edges ratio |
| 0.050 | 0.422 | -0.997 | 0.950 | 0.453 | 0.427 | 0.115 | 0.004 | 0.553 | -1.000 | 1.000 | 0.300 | 0.283 | 0.229 | 0.187 |
| 0.100 | 0.410 | -1.000 | 1.000 | 0.480 | 0.395 | 0.111 | 0.015 | 0.550 | -1.000 | 1.000 | 0.285 | 0.311 | 0.231 | 0.172 |
| 0.150 | 0.396 | -1.000 | 1.000 | 0.507 | 0.357 | 0.110 | 0.026 | 0.548 | -1.000 | 1.000 | 0.274 | 0.336 | 0.236 | 0.155 |
| 0.200 | 0.383 | -1.000 | 1.000 | 0.530 | 0.328 | 0.107 | 0.035 | 0.546 | -1.000 | 1.000 | 0.261 | 0.359 | 0.241 | 0.140 |
| 0.250 | 0.373 | -1.000 | 1.000 | 0.544 | 0.305 | 0.105 | 0.047 | 0.541 | -1.000 | 1.000 | 0.256 | 0.379 | 0.238 | 0.128 |
| 0.300 | 0.364 | -1.000 | 1.000 | 0.551 | 0.280 | 0.107 | 0.062 | 0.539 | -1.000 | 1.000 | 0.253 | 0.391 | 0.240 | 0.116 |
| 0.350 | 0.361 | -1.000 | 1.000 | 0.547 | 0.265 | 0.107 | 0.081 | 0.534 | -1.000 | 1.000 | 0.252 | 0.404 | 0.241 | 0.104 |
| 0.400 | 0.362 | -1.000 | 1.000 | 0.543 | 0.249 | 0.110 | 0.097 | 0.529 | -1.000 | 1.000 | 0.251 | 0.421 | 0.238 | 0.090 |
| 0.450 | 0.351 | -1.000 | 1.000 | 0.552 | 0.236 | 0.109 | 0.103 | 0.525 | -1.000 | 1.000 | 0.254 | 0.439 | 0.228 | 0.079 |
| 0.500 | 0.344 | -1.000 | 1.000 | 0.554 | 0.226 | 0.107 | 0.113 | 0.517 | -1.000 | 1.000 | 0.264 | 0.448 | 0.219 | 0.069 |
| 0.550 | 0.346 | -1.000 | 1.000 | 0.556 | 0.211 | 0.105 | 0.128 | 0.508 | -1.000 | 1.000 | 0.274 | 0.458 | 0.210 | 0.057 |
| 0.600 | 0.314 | -1.000 | 1.000 | 0.573 | 0.197 | 0.104 | 0.127 | 0.500 | -1.000 | 1.000 | 0.281 | 0.471 | 0.201 | 0.047 |
| 0.650 | 0.325 | -1.000 | 1.000 | 0.565 | 0.180 | 0.117 | 0.138 | 0.493 | -1.000 | 1.000 | 0.293 | 0.487 | 0.186 | 0.034 |
| 0.700 | 0.355 | -0.999 | 1.000 | 0.560 | 0.176 | 0.103 | 0.160 | 0.484 | -1.000 | 1.000 | 0.298 | 0.507 | 0.175 | 0.020 |
| 0.750 | 0.330 | -0.998 | 1.000 | 0.561 | 0.190 | 0.097 | 0.151 | 0.477 | -1.000 | 1.000 | 0.310 | 0.517 | 0.162 | 0.011 |
| 0.800 | 0.340 | -1.000 | 0.999 | 0.592 | 0.156 | 0.056 | 0.196 | 0.470 | -0.916 | 0.882 | 0.323 | 0.522 | 0.149 | 0.005 |
| 0.850 | 0.372 | -0.997 | 1.000 | 0.545 | 0.164 | 0.091 | 0.200 | 0.465 | -0.808 | 0.961 | 0.336 | 0.518 | 0.143 | 0.004 |
| 0.900 | 0.450 | -0.652 | 0.999 | 0.444 | 0.333 | 0.000 | 0.222 | 0.461 | -0.341 | 0.859 | 0.347 | 0.510 | 0.139 | 0.003 |
| 0.950 | - | - | - | - | - | - | - | 0.460 | -0.362 | 0.831 | 0.353 | 0.499 | 0.144 | 0.004 |

**Effects of thresholding on ICC in the Brainnetome atlas in unrelated participants.** for each absolute and relative threshold we show the proportion of edges having poor (ICC<0.40), fair (ICC=0.40-0.60), good (ICC=0.60-0.75) or excellent (ICC>0.75) reliability. In this calculation, only subjects for which the edge was retained in both sessions were considered. For absolute thresholds (left) all edges below the value are set to 0, for relative ones (right) only the top percent corresponding to the threshold is retained. A dash indicates that ICC could not be computed for that threshold (not enough remaining consistent edges). Abbreviations: ICC=intraclass correlation coefficient.

Table S13

|  | Absolute | | | | | | | Relative | | | | | | |
| --- | --- | --- | --- | --- | --- | --- | --- | --- | --- | --- | --- | --- | --- | --- |
| Threshold | Median ICC | Min ICC | Max ICC | Poor edges ratio | Fair edges ratio | Good edges ratio | Excellent edges ratio | Median ICC | Min ICC | Max ICC | Poor edges ratio | Fair edges ratio | Good edges ratio | Excellent edges ratio |
| 0.050 | 0.398 | -1.000 | 1.000 | 0.504 | 0.373 | 0.113 | 0.010 | 0.592 | -1.000 | 1.000 | 0.242 | 0.271 | 0.265 | 0.221 |
| 0.100 | 0.360 | -1.000 | 1.000 | 0.559 | 0.303 | 0.109 | 0.029 | 0.583 | -1.000 | 1.000 | 0.239 | 0.296 | 0.276 | 0.189 |
| 0.150 | 0.324 | -1.000 | 1.000 | 0.593 | 0.246 | 0.105 | 0.055 | 0.573 | -1.000 | 1.000 | 0.246 | 0.309 | 0.272 | 0.173 |
| 0.200 | 0.305 | -1.000 | 1.000 | 0.601 | 0.222 | 0.101 | 0.076 | 0.565 | -1.000 | 1.000 | 0.245 | 0.327 | 0.265 | 0.163 |
| 0.250 | 0.290 | -1.000 | 1.000 | 0.608 | 0.203 | 0.099 | 0.089 | 0.559 | -1.000 | 1.000 | 0.245 | 0.343 | 0.260 | 0.152 |
| 0.300 | 0.276 | -1.000 | 1.000 | 0.616 | 0.189 | 0.097 | 0.099 | 0.554 | -1.000 | 1.000 | 0.245 | 0.358 | 0.255 | 0.143 |
| 0.350 | 0.266 | -1.000 | 1.000 | 0.614 | 0.185 | 0.095 | 0.107 | 0.547 | -1.000 | 1.000 | 0.246 | 0.377 | 0.247 | 0.131 |
| 0.400 | 0.277 | -1.000 | 1.000 | 0.601 | 0.177 | 0.096 | 0.127 | 0.542 | -1.000 | 1.000 | 0.247 | 0.388 | 0.247 | 0.118 |
| 0.450 | 0.286 | -1.000 | 1.000 | 0.597 | 0.171 | 0.093 | 0.139 | 0.534 | -1.000 | 1.000 | 0.250 | 0.407 | 0.241 | 0.102 |
| 0.500 | 0.300 | -1.000 | 1.000 | 0.583 | 0.164 | 0.111 | 0.142 | 0.527 | -1.000 | 1.000 | 0.253 | 0.428 | 0.235 | 0.084 |
| 0.550 | 0.278 | -1.000 | 1.000 | 0.584 | 0.158 | 0.106 | 0.151 | 0.519 | -1.000 | 1.000 | 0.259 | 0.449 | 0.226 | 0.065 |
| 0.600 | 0.334 | -1.000 | 1.000 | 0.559 | 0.195 | 0.104 | 0.142 | 0.510 | -1.000 | 1.000 | 0.265 | 0.471 | 0.217 | 0.046 |
| 0.650 | 0.340 | -0.998 | 1.000 | 0.561 | 0.203 | 0.114 | 0.122 | 0.501 | -1.000 | 1.000 | 0.271 | 0.495 | 0.204 | 0.031 |
| 0.700 | 0.328 | -0.969 | 0.945 | 0.593 | 0.204 | 0.106 | 0.097 | 0.493 | -1.000 | 0.999 | 0.278 | 0.516 | 0.187 | 0.019 |
| 0.750 | 0.372 | -1.000 | 0.967 | 0.619 | 0.190 | 0.079 | 0.111 | 0.485 | -0.994 | 1.000 | 0.289 | 0.530 | 0.170 | 0.010 |
| 0.800 | 0.056 | -1.000 | 0.976 | 0.655 | 0.138 | 0.138 | 0.069 | 0.477 | -0.796 | 0.998 | 0.302 | 0.538 | 0.154 | 0.006 |
| 0.850 | 0.358 | -0.674 | 0.909 | 0.500 | 0.333 | 0.000 | 0.167 | 0.469 | -0.796 | 0.981 | 0.319 | 0.535 | 0.142 | 0.004 |
| 0.900 | 0.786 | 0.786 | 0.786 | 0.000 | 0.000 | 0.000 | 1.000 | 0.460 | -0.416 | 0.921 | 0.339 | 0.524 | 0.133 | 0.003 |
| 0.950 | - | - | - | - | - | - | - | 0.455 | -0.349 | 0.870 | 0.356 | 0.512 | 0.129 | 0.003 |

**Effects of thresholding on ICC in the Glasser atlas in unrelated participants.** for each absolute and relative threshold we show the proportion of edges having poor (ICC<0.40), fair (ICC=0.40-0.60), good (ICC=0.60-0.75) or excellent (ICC>0.75) reliability. In this calculation, only subjects for which the edge was retained in both sessions were considered. For absolute thresholds (left) all edges below the value are set to 0, for relative ones (right) only the top percent corresponding to the threshold is retained. A dash indicates that ICC could not be computed for that threshold (not enough remaining consistent edges). Abbreviations: ICC=intraclass correlation coefficient.

Table S14

|  | Absolute | | | | | | | Relative | | | | | | |
| --- | --- | --- | --- | --- | --- | --- | --- | --- | --- | --- | --- | --- | --- | --- |
| Threshold | Median ICC | Min ICC | Max ICC | Poor edges ratio | Fair edges ratio | Good edges ratio | Excellent edges ratio | Median ICC | Min ICC | Max ICC | Poor edges ratio | Fair edges ratio | Good edges ratio | Excellent edges ratio |
| 0.050 | 0.363 | -1.000 | 1.000 | 0.561 | 0.327 | 0.095 | 0.017 | 0.572 | -1.000 | 1.000 | 0.276 | 0.273 | 0.243 | 0.208 |
| 0.100 | 0.329 | -1.000 | 1.000 | 0.596 | 0.268 | 0.094 | 0.042 | 0.560 | -1.000 | 1.000 | 0.282 | 0.292 | 0.241 | 0.184 |
| 0.150 | 0.300 | -1.000 | 1.000 | 0.612 | 0.231 | 0.094 | 0.062 | 0.549 | -1.000 | 1.000 | 0.281 | 0.319 | 0.239 | 0.161 |
| 0.200 | 0.281 | -1.000 | 1.000 | 0.622 | 0.202 | 0.093 | 0.083 | 0.540 | -1.000 | 1.000 | 0.284 | 0.335 | 0.232 | 0.149 |
| 0.250 | 0.258 | -1.000 | 1.000 | 0.628 | 0.179 | 0.095 | 0.097 | 0.533 | -1.000 | 1.000 | 0.285 | 0.354 | 0.222 | 0.139 |
| 0.300 | 0.251 | -1.000 | 1.000 | 0.627 | 0.174 | 0.090 | 0.109 | 0.526 | -1.000 | 1.000 | 0.292 | 0.363 | 0.217 | 0.129 |
| 0.350 | 0.217 | -1.000 | 1.000 | 0.636 | 0.160 | 0.092 | 0.112 | 0.517 | -1.000 | 1.000 | 0.299 | 0.375 | 0.208 | 0.118 |
| 0.400 | 0.237 | -1.000 | 1.000 | 0.622 | 0.164 | 0.091 | 0.122 | 0.508 | -1.000 | 1.000 | 0.310 | 0.385 | 0.204 | 0.102 |
| 0.450 | 0.262 | -1.000 | 1.000 | 0.620 | 0.167 | 0.094 | 0.119 | 0.499 | -1.000 | 1.000 | 0.321 | 0.398 | 0.196 | 0.085 |
| 0.500 | 0.331 | -0.999 | 1.000 | 0.563 | 0.207 | 0.099 | 0.131 | 0.489 | -1.000 | 1.000 | 0.332 | 0.416 | 0.186 | 0.067 |
| 0.550 | 0.318 | -0.997 | 1.000 | 0.556 | 0.183 | 0.114 | 0.147 | 0.478 | -1.000 | 1.000 | 0.342 | 0.436 | 0.176 | 0.046 |
| 0.600 | 0.383 | -1.000 | 1.000 | 0.514 | 0.229 | 0.107 | 0.150 | 0.468 | -1.000 | 1.000 | 0.355 | 0.454 | 0.163 | 0.028 |
| 0.650 | 0.350 | -0.990 | 1.000 | 0.571 | 0.180 | 0.068 | 0.180 | 0.459 | -0.999 | 1.000 | 0.369 | 0.469 | 0.147 | 0.015 |
| 0.700 | 0.407 | -0.975 | 0.979 | 0.491 | 0.218 | 0.145 | 0.145 | 0.449 | -0.971 | 0.982 | 0.386 | 0.476 | 0.130 | 0.008 |
| 0.750 | -0.057 | -0.962 | 0.934 | 0.625 | 0.208 | 0.000 | 0.167 | 0.440 | -0.732 | 0.902 | 0.407 | 0.474 | 0.115 | 0.004 |
| 0.800 | 0.074 | -0.853 | 0.557 | 0.750 | 0.250 | 0.000 | 0.000 | 0.429 | -0.643 | 0.855 | 0.429 | 0.465 | 0.103 | 0.003 |
| 0.850 | 0.517 | 0.122 | 0.971 | 0.333 | 0.333 | 0.000 | 0.333 | 0.420 | -0.464 | 0.825 | 0.452 | 0.451 | 0.095 | 0.002 |
| 0.900 | 0.844 | 0.844 | 0.844 | 0.000 | 0.000 | 0.000 | 1.000 | 0.413 | -0.346 | 0.825 | 0.470 | 0.436 | 0.091 | 0.002 |
| 0.950 | - | - | - | - | - | - | - | 0.407 | -0.305 | 0.825 | 0.483 | 0.425 | 0.090 | 0.002 |

**Effects of thresholding on ICC in the Gordon atlas in unrelated participants.** for each absolute and relative threshold we show the proportion of edges having poor (ICC<0.40), fair (ICC=0.40-0.60), good (ICC=0.60-0.75) or excellent (ICC>0.75) reliability. In this calculation, only subjects for which the edge was retained in both sessions were considered. For absolute thresholds (left) all edges below the value are set to 0, for relative ones (right) only the top percent corresponding to the threshold is retained. A dash indicates that ICC could not be computed for that threshold (not enough remaining consistent edges). Abbreviations: ICC=intraclass correlation coefficient.

**Supplementary Figures**

Figure S1

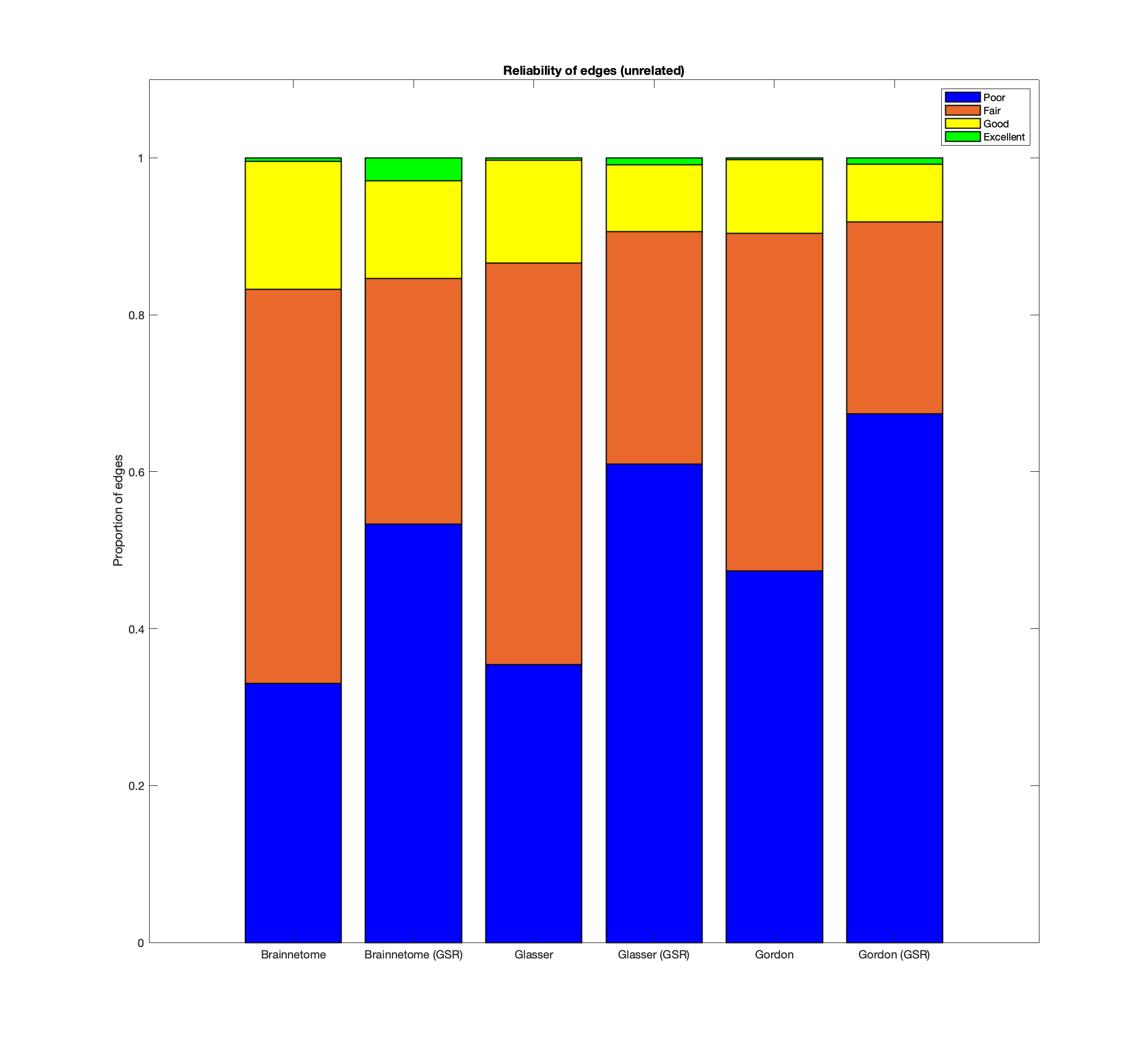

**Reliability of functional connectome edges in unrelated participants.** For the Brainnetome, Glasser and Gordon atlases with and without performing GSR we show the proportion of edges having poor (ICC<0.40), fair (ICC=0.40-0.60), good (ICC=0.60-0.75) or excellent (ICC>0.75) reliability, defined in accordance to ^20^. Abbreviations: ICC=intraclass correlation coefficient, GSR-=no global signal regression, GSR+=global signal regression.

Figure S2

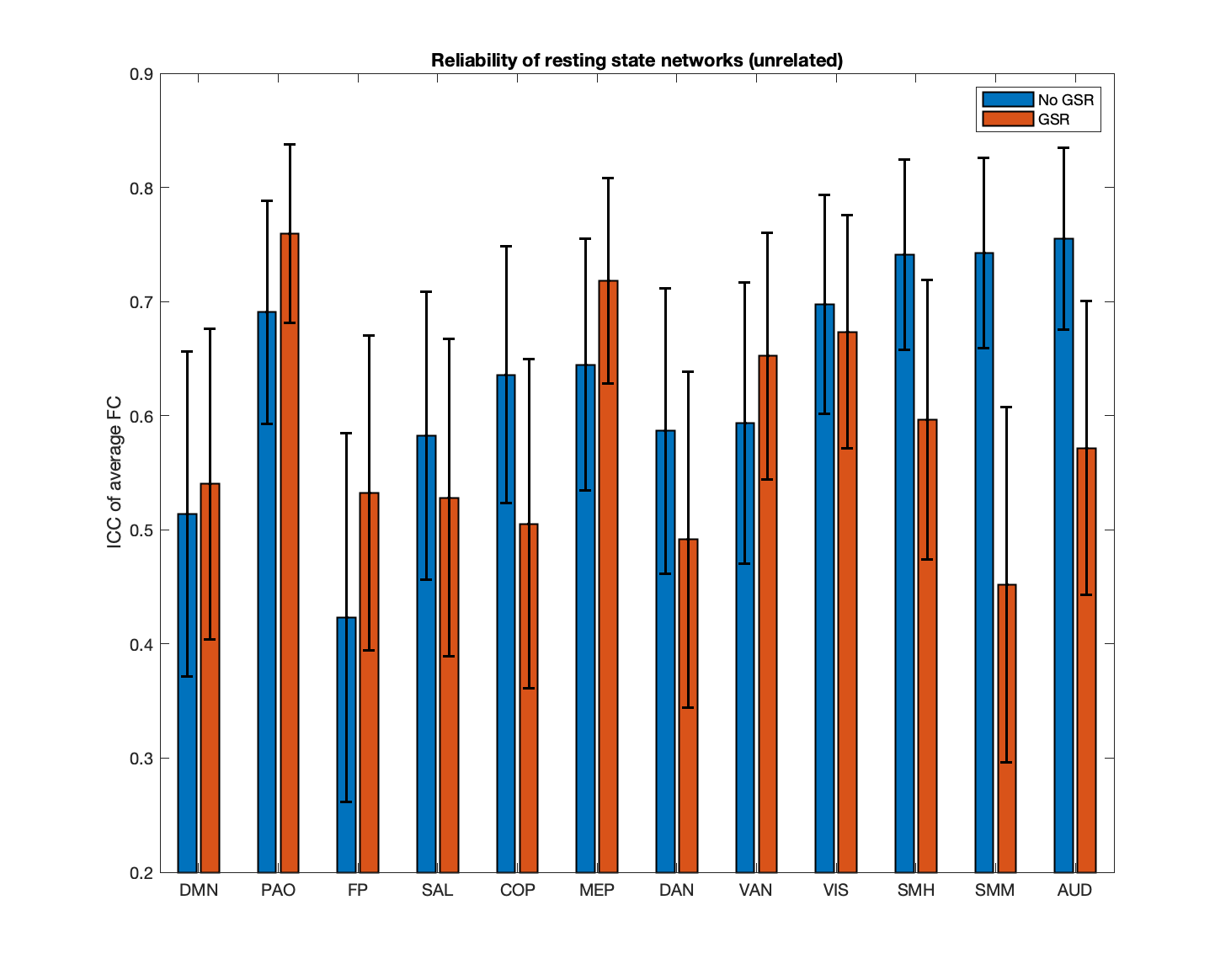

**Reliability of known resting state networks in unrelated participants.** We show the ICC and confidence intervals for the average connectivity within known resting state networks defined in accordance to ^28^. Abbreviations: ICC=intraclass correlation coefficient, GSR-=no global signal regression, GSR+=global signal regression, CI=confidence interval, DMN=default mode network, PAO=parieto-occipital, FP=fronto-parietal, SAL=salience, COP=cingulo-opercular, MEP=medial parietal, DAN=dorsal attention network, VAN=ventral attention network, VIS=visual, SMH=supplementary motor (hand), SMM=supplementary motor (mouth), AUD=auditory.

Figure S3

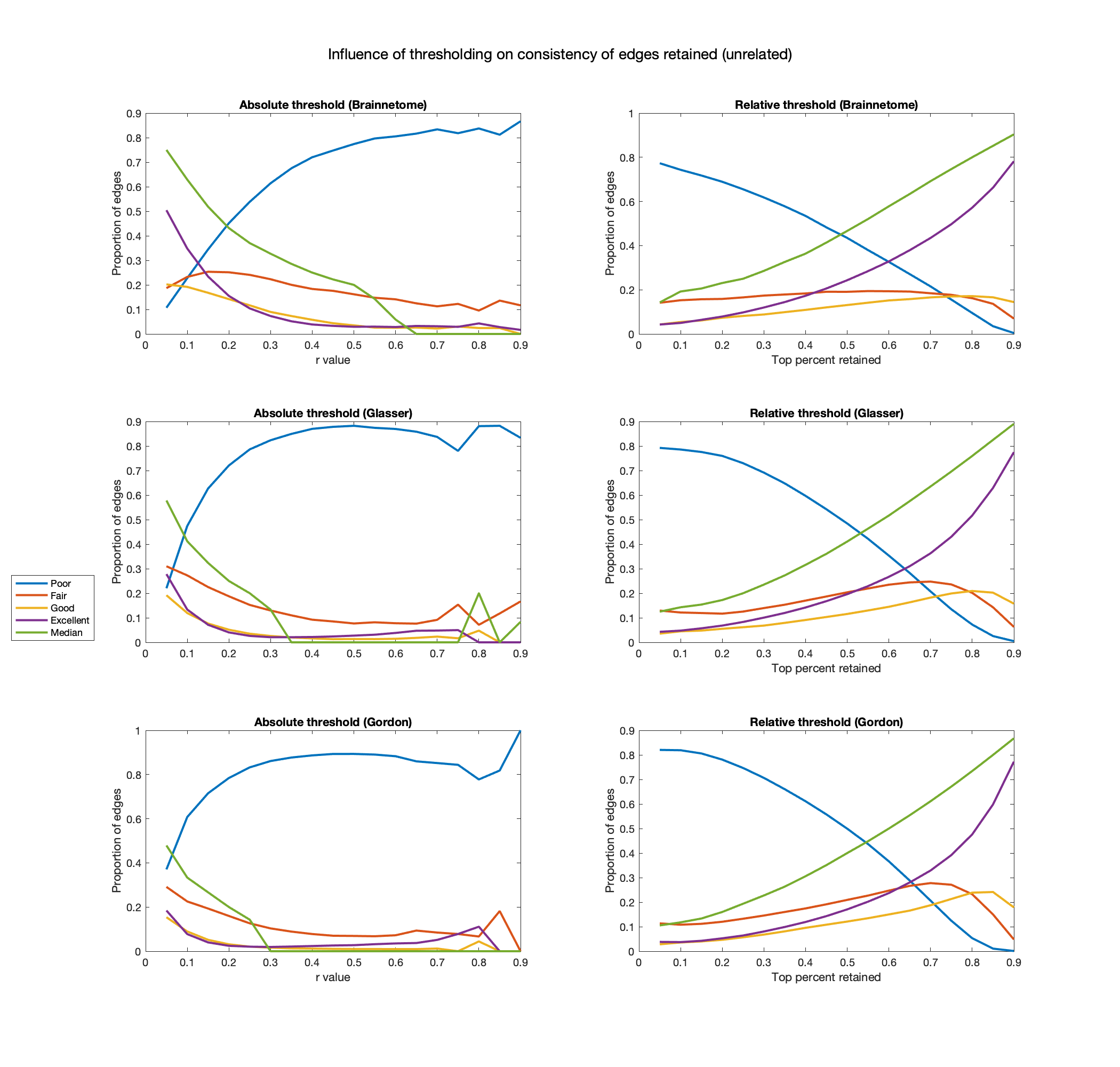

**Effects of thresholding on edge retention in unrelated participants.** In the Brainnetome, Glasser and Gordon atlases, for each absolute and relative threshold we show the proportion of edges that are consistently retained. As a measure of consistency, we use the number of participants in which the edge was retained at both timepoints divided by the ones in which it was retained at least once. For convenience, we then use the values defined in ^20^ to plot the ratio of edges having poor (ratio<0.40), fair (ratio=0.40-0.60), good (ratio=0.60-0.75) or excellent (ratio >0.75) consistency. For absolute thresholds (left) all edges below the value are set to 0, for relative ones (right) only the top percent corresponding to the threshold is retained. Abbreviations: r=Pearson correlation coefficient.

Figure S4

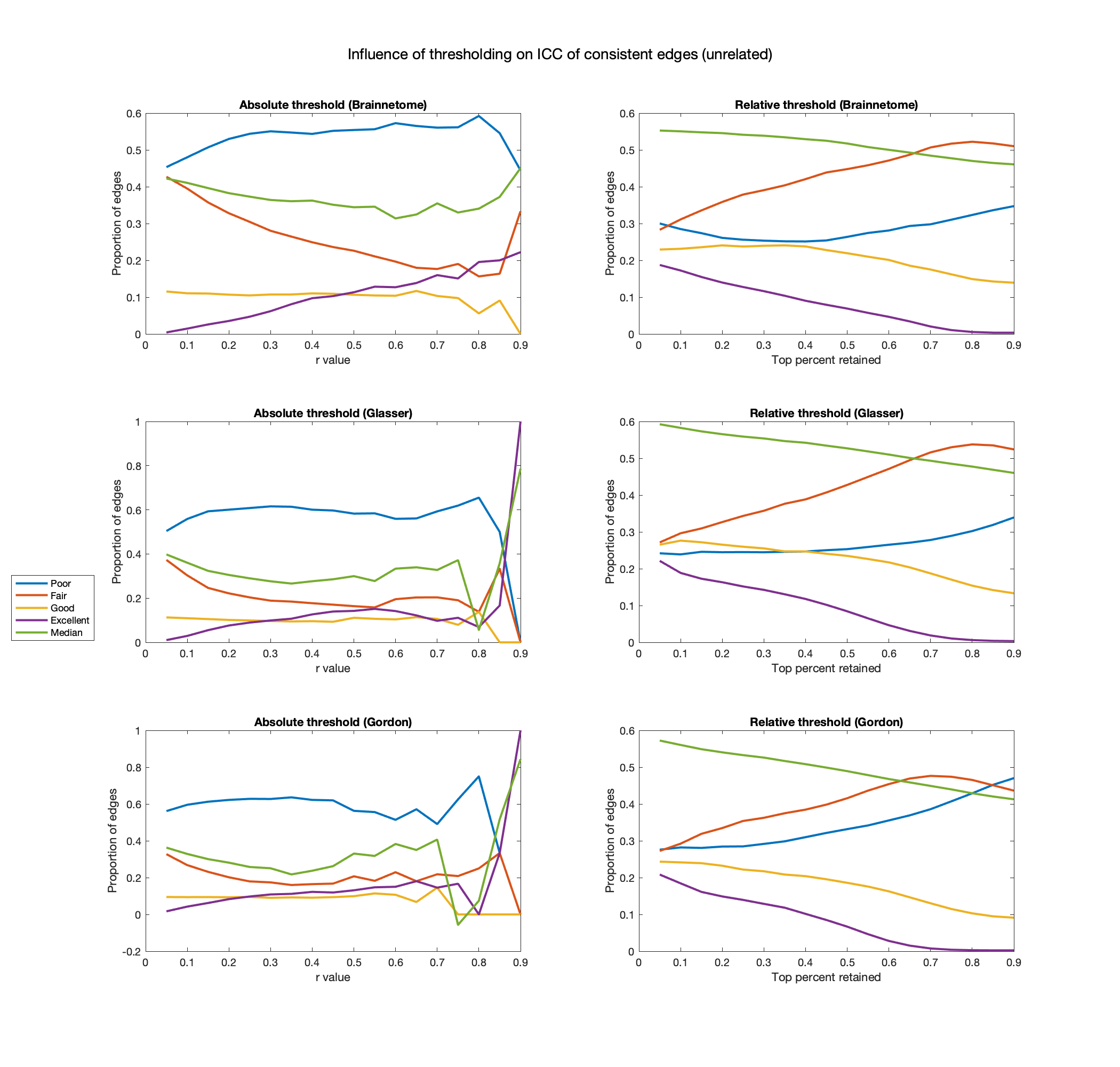

**Effects of thresholding on reliability in unrelated participants.** In the Brainnetome, Glasser and Gordon atlases, for each absolute and relative threshold we show the proportion of edges having poor (ICC<0.40), fair (ICC=0.40-0.60), good (ICC=0.60-0.75) or excellent (ICC>0.75) reliability. In this calculation, only subjects for which the edge was retained in both sessions were considered. For absolute thresholds (left) all edges below the value are set to 0, for relative ones (right) only the top percent corresponding to the threshold is retained. Abbreviations: r=Pearson correlation coefficient, ICC=intraclass correlation coefficient.
